# Structures of LIG1 engaging with mutagenic mismatches inserted by polβ in base excision repair

**DOI:** 10.1101/2022.01.14.473406

**Authors:** Qun Tang, Robert McKenna, Melike Çağlayan

## Abstract

DNA ligase I (LIG1) catalyzes final ligation step following DNA polymerase (pol) β gap filling and an incorrect nucleotide insertion by polβ creates a nick repair intermediate with mismatched end at the downstream steps of base excision repair (BER) pathway. Yet, how LIG1 discriminates against the mutagenic 3′-mismatches at atomic resolution remains undefined. Here, we determined X-ray structures of LIG1/nick DNA complexes with G:T and A:C mismatches and uncovered the ligase strategies that favor or deter ligation of base substitution errors. Our structures revealed that LIG1 active site can accommodate G:T mismatch in a similar conformation with A:T base pairing, while it stays in the LIG1-adenylate intermediate during initial step of ligation reaction in the presence of A:C mismatch at 3′-strand. Moreover, we showed mutagenic ligation and aberrant nick sealing of the nick DNA substrates with 3′-preinserted dG:T and dA:C mismatches, respectively. Finally, we demonstrated that AP-Endonuclease 1 (APE1), as a compensatory proofreading enzyme, interacts and coordinates with LIG1 during mismatch removal and DNA ligation. Our overall findings and ligase/nick DNA structures provide the features of accurate versus mutagenic outcomes at the final BER steps where a multi-protein complex including polβ, LIG1, and APE1 can maintain accurate repair.

## INTRODUCTION

DNA strand breaks occur as intermediates during all DNA transactions including DNA replication, DNA repair, and genetic recombination (1). Human DNA ligases catalyze a phosphodiester bond formation between 5′-phosphate (P) and 3′-hydroxyl (OH) termini on ends of the broken DNA strand, and therefore contribute to overall genome stability (2–5). Using a high-energy cofactor ATP and Mg^2+^, DNA ligases catalyze the ligation reaction including three chemical and sequential steps: (i) nucleophilic attack on ATP by the ligase and formation of a covalent intermediate in which adenylate (AMP) is linked to an active site lysine (LIG-AMP), (ii) the AMP is transferred to the 5′-end of the 5′-phosphate-terminated DNA strand to form DNA-AMP intermediate, and (iii) the ligase catalyzes attack by 3′-OH of the nick on DNA-adenylate to join adjacent 3′-OH and 5′-P ends and liberate AMP (6–13).

The accuracy of the nick sealing reaction at the end of DNA repair relies on the formation of a Watson-Crick base pair between the 5′-P and 3′-OH ends that requires high fidelity DNA synthesis by DNA polymerase (14). Base excision repair (BER) is the predominant DNA repair mechanism of small single-base DNA lesions (15). The BER pathway involves the mechanism of substrate-product channeling that entails subsequent enzymatic steps and hand off of the repair intermediates between BER proteins so that the release and accumulation of toxic and mutagenic single-strand break intermediates are minimized in cells (16–20). This repair pathway coordination involves consecutive DNA synthesis and nick sealing steps during which DNA polymerase (pol) β incorporates a single nucleotide into a gap and the resulting nicked insertion product is handed off to the ligation step where DNA ligase (ligase I and IIIα) joins 3′-OH and 5′-P ends to complete the BER pathway (14,21). Polβ, an error-prone polymerase without 3′-5′ exonuclease proofreading activity, can incorporate mismatch nucleotides at a frequency of 1 in ~5000 during template-directed DNA synthesis (22). This polβ mismatch product could generate a problematic nick repair intermediate for subsequent ligation step in the BER pathway (14,21). In the presence of 3′-damaged or modified DNA ends, DNA ligases can fail resulting in the formation of 5′-adenylated-DNA intermediates (*i.e*. 5′-AMP), also referred to abortive ligation products (23–25).

In our previous studies, we demonstrated the importance of polβ and DNA ligase I (LIG1) coordination for the accurate channeling of repair intermediates to maintain BER efficiency at the downstream steps of the repair pathway (23–36). For example, we reported that the incorporation of oxidized nucleotide 7,8-dihydro-8′-oxo-dGTP (8-oxodGTP) by polβ confounds LIG1, leading to the formation of abortive repair intermediates (26,27). In our recent studies, we demonstrated that the formation of polβ correct nucleotide insertion product that shows a stable closed ternary complex conformation enables the recognition and ligation of the nicked insertion product and its efficient hand off to next ligation step in the BER pathway (32). Furthermore, we showed that the repair fidelity is affected by the situations involving 5-methylcytosine (5mC) and oxidative 5mC base modifications in template DNA and the disease-associated mutations linked to LIG1-deficiency syndrome lead to enhanced ligation failure after polβ 8-oxodGTP insertion (33,34). Recently, we demonstrated the mechanism by which LIG1 fidelity mediates the faithful substrate-product channeling, specifically low-fidelity LIG1 results in the mutagenic ligation of 8-oxodGMP inserted by polβ (35). These studies contribute to understanding of important molecular determinants that ensure accurate BER pathway coordination or result in impaired hand off from polβ to DNA ligase. However, it still remains unclear that how DNA ligase dictates accurate versus mutagenic outcomes for polβ-mediated base substitution errors during the final nick sealing step of BER pathway.

Polβ has been reported to be mutated in 30% of a variety of human tumors such as lung, gastric, colorectal, and prostate cancer (37,38). Several of the cancer-associated polβ variants possess aberrant repair function *in vitro* such as a reduced fidelity stemming from impaired discrimination against incorrect nucleotide incorporation as reported for polβ cancer-associated mutant K289M (39,40). The expression of these variants in cells induces cellular transformation and genomic instability (41–46). The mismatch nucleotide insertions by DNA polymerases during repair and replication processes can cause base substitutions, additions and deletions in case of no proofreading, leading to genome instability and human diseases (47). For example, G:T mismatches are among the more prevalent mismatches found in nature, arising from the deamination of 5-methylcytosine to thymine (48,49). Furthermore, it has been reported that Watson-Crick like G:T mismatch, if left unrepaired, could lead to transition or transversion point mutations and be a prominent source of base substitution errors in tumor suppressor genes in multiple forms of cancer (50–53). Nevertheless, the extent to which discrimination by LIG1 counteracts mutagenic polβ mismatch insertion-promoted repair intermediates at atomic resolution remains unknown.

In the present study, we defined the molecular basis of human LIG1 mismatch discrimination mechanism by moderate-resolution structures of LIG1/nick DNA duplexes harboring A:C and G:T mismatches at 3′-end. Our structures revealed that LIG1 active site can accommodate G:T mismatch in a similar conformation with A:T base pairing where the ligase validates mutagenic G:T ligation during the adenyl transfer step of ligation reaction (DNA-AMP). However, the ligase active site exhibits large distortion where the position of 3′-OH end rotates 50° from the nick DNA with A:C mismatch during the first step of ligation reaction where the active site lysine (K568) residue stays adenylated (LIG1-AMP) and 5′-PO_4_ end shows a conformational change. In our ligation assays *in vitro*, we showed efficient and defective nick sealing of G:T and A:C mismatches, respectively. Furthermore, we found that LIG1 can ligate the nicked repair product following polβ dGTP:T mismatch insertion and deters nick sealing after dATP:C insertion. Finally, our results demonstrated that, APE1, as a complementary proofreading enzyme, can remove a mismatched base from 3′-end of the nick DNA substrates by its exonuclease activity. We also showed APE1/LIG1 protein-protein interaction and functional coordination for mismatch removal coupled to DNA ligation. Overall results reveal the strategies of LIG1 engaging with mismatched nick DNA that govern ligation of base substitution errors inserted by polβ and demonstrate the requirement of a multi-protein assembly (polβ, LIG1, and APE1) to maintain the repair efficiency at the downstream steps of the BER pathway.

## RESULTS

### LIG1 engaging with nick repair intermediates with mismatched DNA ends

In our previous study, we reported that the polβ mismatch insertion governs the channeling of resulting nicked repair product to LIG1 with the exception of Watson-Crick-like dGTP insertion opposite T (32). In order to elucidate the ligase strategies that deter or favor the ligation of repair intermediates that mimic polβ mismatch nucleotide insertion products at atomic resolution, we determined the structures of LIG1/nick DNA duplexes with G:T and A:C mismatches and Watson-Crick A:T base-pair (Table 1).

**Table 1.**
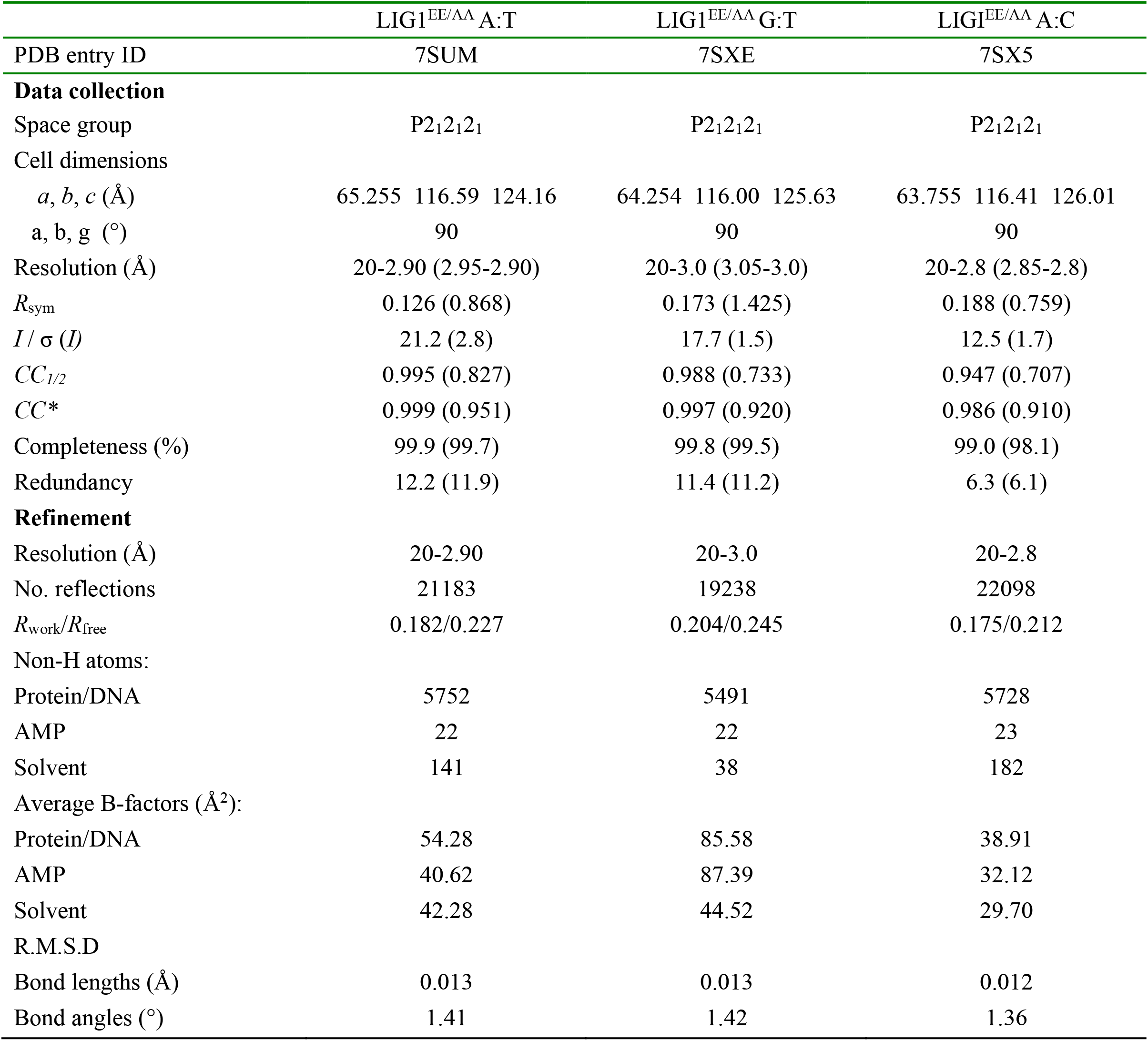
X-ray data collection and refinement statistics of LIG1/nick DNA duplexes with A:T, G:T and A:C ends at 3′-strand.

Our structures demonstrated the molecular mechanism of LIG1 engaging with nick DNA harboring mutagenic mismatch at 3′-strand (Figure 1 and Supplementary Figure 1). In the structure of LIG1 bound to nick DNA duplexes containing A:T and G:T, we showed that the ligase active site can accommodate G:T mismatch in a similar conformation with Watson-Crick A:T base pairing (Figure 1A and 1B). The structure comparisons revealed no significant differences, with superimposition Cα root mean square deviation of 0.609 Å. In both LIG1/nick DNA structures, the 5′-termini is adenylated in the crystals and DNA-adenylate (DNA-AMP) intermediate is observed. This refers to the second step of ligation reaction when AMP is transferred to 5′-PO_4_ of nick DNA and shows that LIG1/nick conformation with A:T and G:T is poised for step 3 (nick sealing).

**Figure 1.**
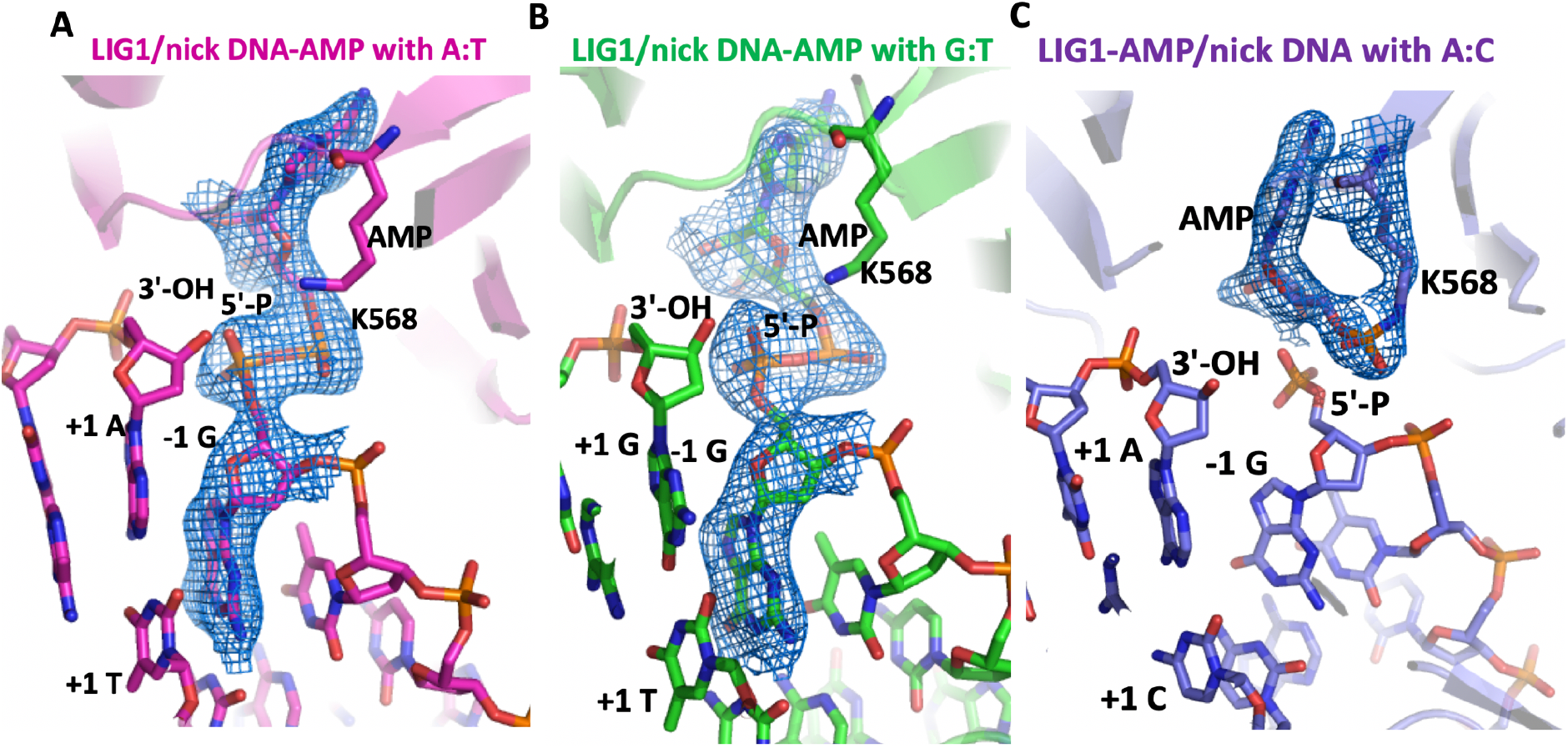
Structures of LIG1 bound to nick DNA duplexes with G:T and A:C mismatches. X-ray crystal structures of LIG1/nick DNA duplexes with A:T (**A**), G:T (**B**), and A:C (**C**) at 3′-strand. LIG1/A:T (magenta) and G:T (green) structures with AMP-DNA complex weighted 2Fo-Fc electron density contoured at 1σ are displayed for the adenylated 5′-phosphate of the nick (DNA-AMP). The structure of LIG1/A:C (blue) with the ligase-AMP complex weighted 2Fo-Fc electron density contoured at 1σ is displayed for adenylated LIG1 (LIG1-AMP) at K568 active site residue. DNA and LIG1 are shown as sticks and cartoon, respectively, and AMP is depicted in blue.

In contrast, significant rearrangements at LIG1 active site were observed near the A:C mismatch. The structure of LIG1/nick DNA duplex with A:C revealed that the ligase active site exhibits LIG1-adenylate conformation where the active side lysine residue (K568) is covalently bound to the AMP phosphate (Figure 1C). This refers to the first step of ligation reaction and indicates that LIG1 stays in its initial adenylated state and cannot move forward with subsequent adenyl group transfer to the 5′-PO_4_ on the downstream strand to activate the ligase for attack by the upstream 3′-OH of nick DNA. Our LIG1/A:C mismatch structure represents the first human LIG1 structure resolved in step 1. These observations suggest that the A:C base pairing imparts non-native active site conformations that further suppress the chemical steps of catalysis. Overall, our LIG1/mismatch structures demonstrate that the ligase is trapped as the adenylated-DNA intermediate (AMP-DNA) that favors the ligation of mutagenic G:T mismatch and the active site remains in inactive conformation (AMP-K568) that deters nick sealing of A:C end.

Similar to the previously reported LIG1 structures (54–57), we observed that LIG1 completely envelopes the DNA with DNA binding (DBD), adenylylation (AdD), and oligonucleotide binding (OBD) domains encircling the nick DNA with correct or mismatched ends (Supplementary Figure 2). Moreover, we observed that the structures of LIG1/A:T nick DNA harbors a Watson-Crick conformation with two hydrogen bonds, while G:T and A:C 3′-terminal pairs show the Wobble hydrogen bonding, which is similar to DNA polymerase/mismatch structures for G:T and A:C (Figure 2 and Supplementary Figure 3).

**Figure 2.**
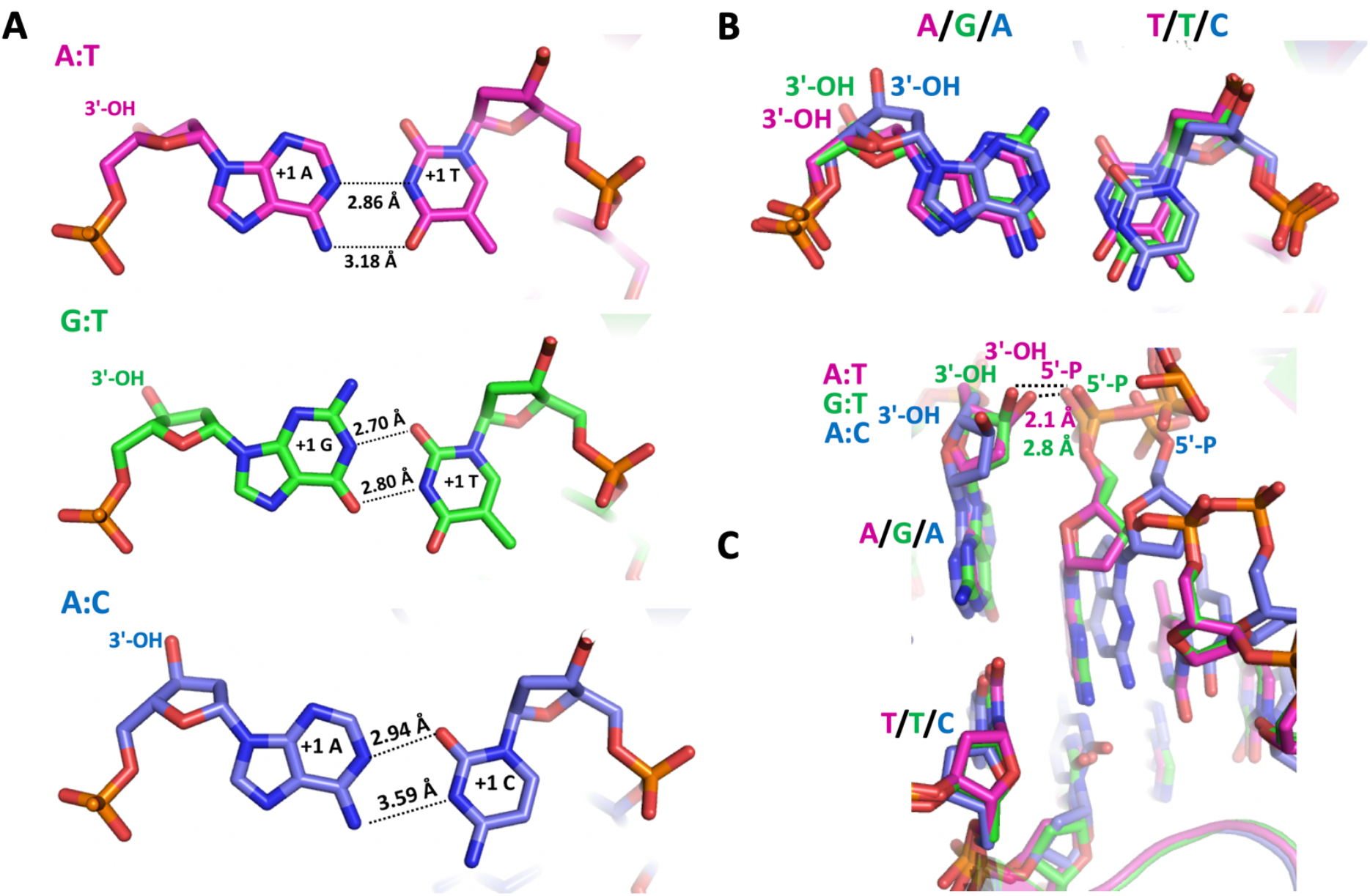
LIG1/nick duplexes with mismatched ends exhibit different DNA conformations. (**A**) Hydrogen-bonding patterns of A:T, G:T, and A:C base-pairs. (**B,C**) Overlay of LIG1 X-ray structures bound to the nick DNA duplexes with A:T (magenta), G:T (green) and A:C (blue). The mismatched 3′-OH strand bound in the LIG1/nick DNA complexes (B) is depicted to show differences in the distances between 5′-P and 3′-OH ends of a nick (C).

### LIG1 active site shows distinct DNA conformations depending on the identity of mismatched ends

High-fidelity DNA synthesis requires that the polymerases display a strong preference for right nucleotide insertion (22). Previously solved polβ/mismatch structures indicated that the mismatched termini adopt various distorted conformations that attempt to satisfy stacking and hydrogen-bonding interactions, which provides a key fidelity checkpoint (58–68). Our LIG1 structures in complex with nick DNA harboring G:T and A:C mismatched termini exhibit distinct mismatch-specific conformations. We found significant differences in the position of 5′-PO_4_ and 3′-OH strands at a nick around the upstream and downstream DNA (Figures 2–5). For the structures of LIG1 in complex with nick DNA containing G:T and A:T where the 5′-5′ phosphoanhydride AMP-DNA intermediate is formed, we observed that 5′-phosphate is more close to 3′-OH strand of a nick for proper positioning and sealing a phosphodiester backbone (Figure 2C). The distances from 5′-P to 3′-OH of nick DNA with A:T and G:T are 2.1 Å and 2.8 Å, respectively. In LIG1/A:C mismatch structure, we observed that 3′-OH of nick DNA rotates 50° from that of nick DNA with A:T (Figure 2C). The overlay of LIG1/nick structures with correctly base-paired A:T versus mismatched G:T or A:C ends also demonstrated significant differences in the conformations of 5′-strand (Figure 3A), due to clear shifts in the positions of -1G, -2T, and -3C nucleotides relative of the upstream DNA in the structures of LIG1/G:T (Figure 3A) and A:C (Figure 3B) mismatches in comparison with LIG1/A:T nick DNA (Supplementary Figure 1).

**Figure 3.**
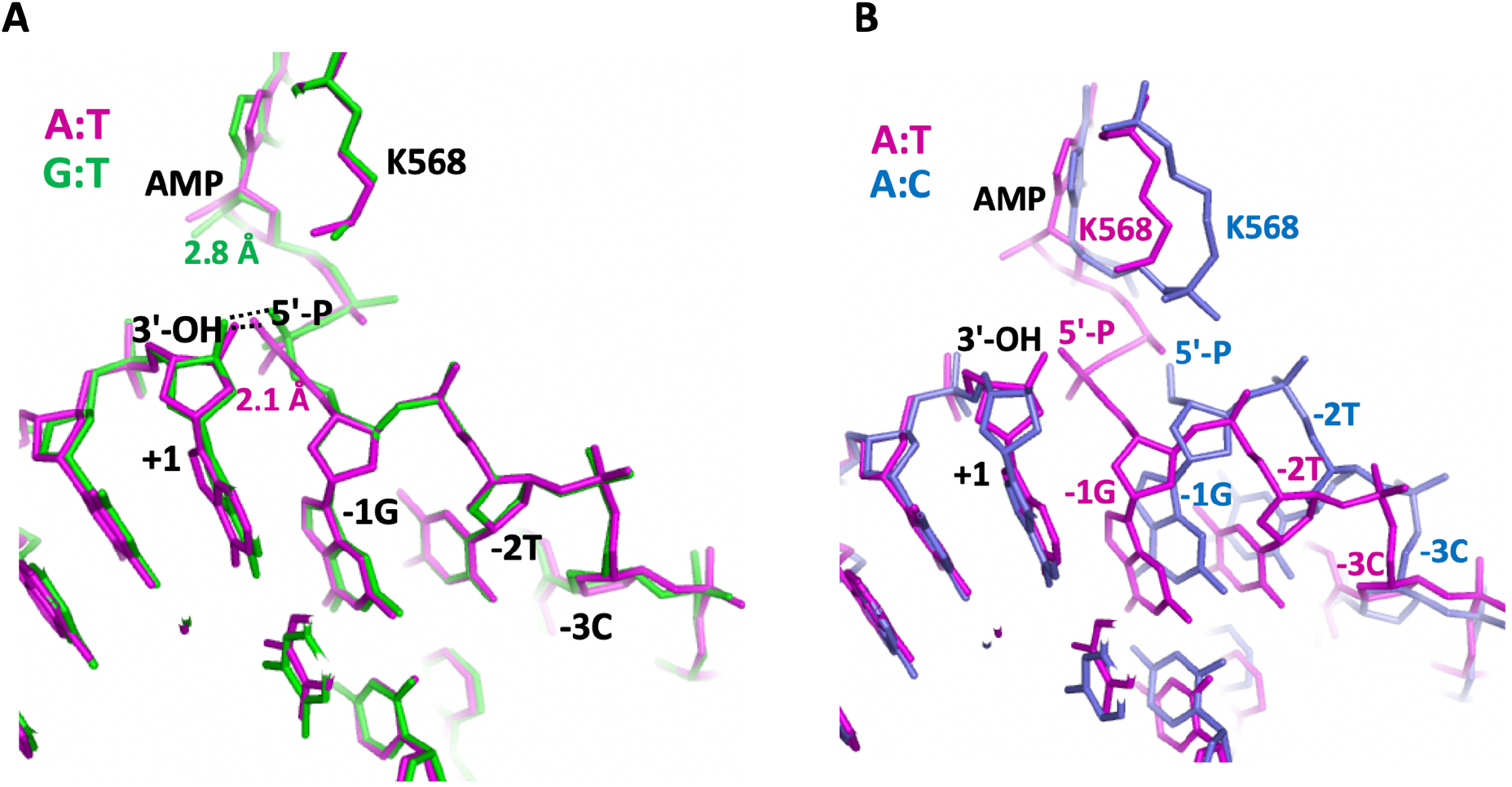
LIG1 active site bound to nick DNA duplexes with mismatched ends. Overlay of LIG1/nick DNA duplexes for A:T/G:T (magenta/green) and A:T/A:C (magenta/blue) structures in panels A and B, respectively. The superimposition of LIG1/nick duplexes show the differences in the position of adenylate (AMP) bound to nick DNA (G:T mismatch) or K568 active site (A:C mismatch) of LIG1.

Moreover, we observed the position of phenylalanine at 872 (F872) that is located upstream of the nick and positioned close to the deoxyribose moiety of the nucleotide at the 5′-end shows differences in the LIG1/mismatch nick DNA structures (Figure 4A). The overlay of LIG1/nick DNA duplexes with A:T and A:C demonstrated that F872 distorts the alignment at the upstream of A:T nick where -1G and +1A nucleotides are in parallel between 3′- and 5′-strands (Figure 4B-C). In both LIG1 structures, we also found conformational differences in the active site residues Arg(R)589 and Leu(L)544, which are positioned close to 5′-phosphate of a nick (Figure 5). The interaction interface between Arg(R)874 and -2T nucleotide of the downstream DNA shows a clear change in LIG1 structures with A:T versus A:C ends (Figure 5A). Similarly, the distance between R589 and L544 side chains is shifted because of the differences in the position of AMP (bound to DNA at LIG1/A:T or bound to K568 active site at LIG1/A:C) as shown in the LIG1 A:T/A:C overlay structure (Figure 5B).

**Figure 4.**
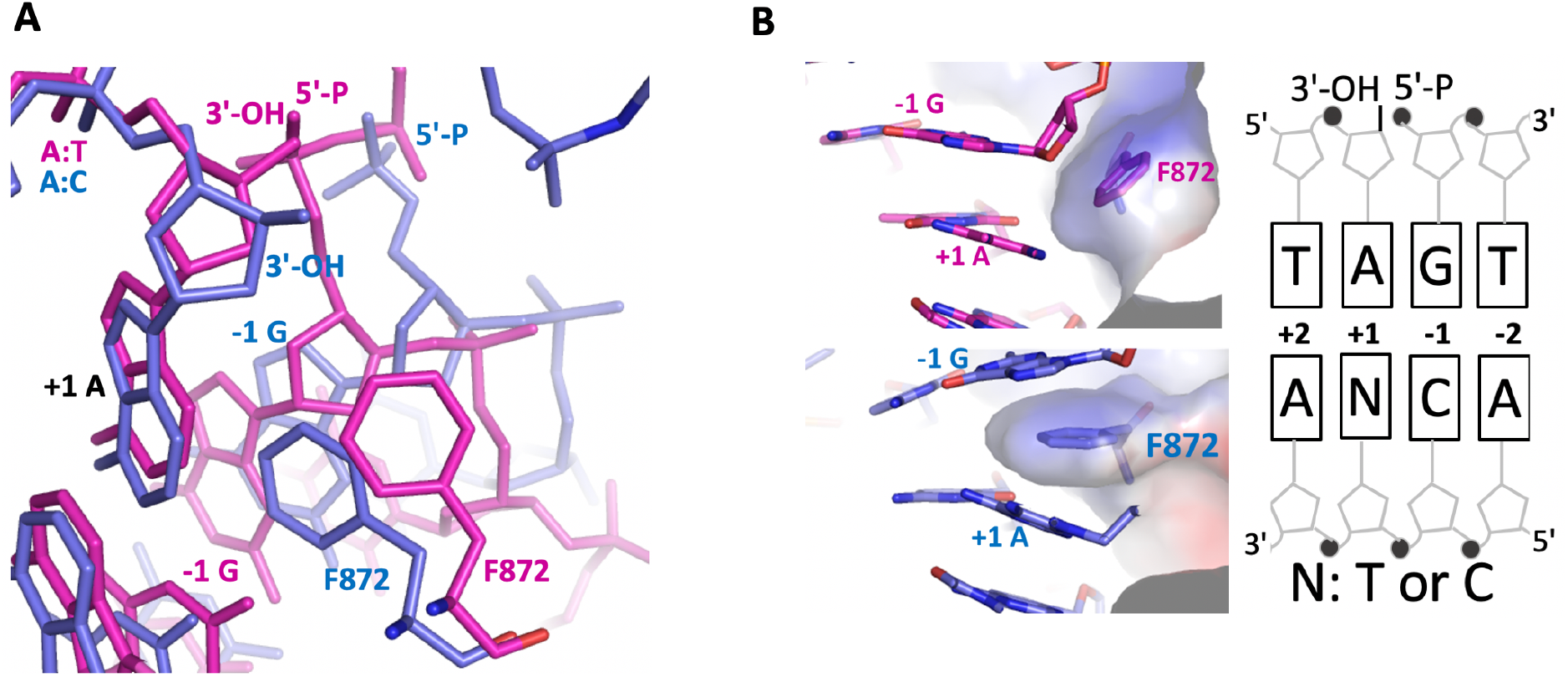
Structures of LIG1 F872 in complex with nick DNA complexes with mismatched ends. **(A)** Overlay of LIG1 X-ray structures bound to the nick DNA duplexes with A:T (magenta) and A:C (blue) show the conformational change at F872 (stick). (**B**) Surface representations of LIG1 X-ray structures depict protein-DNA contacts at F872 with +1A and - 1G bases of nick DNA.

**Figure 5.**
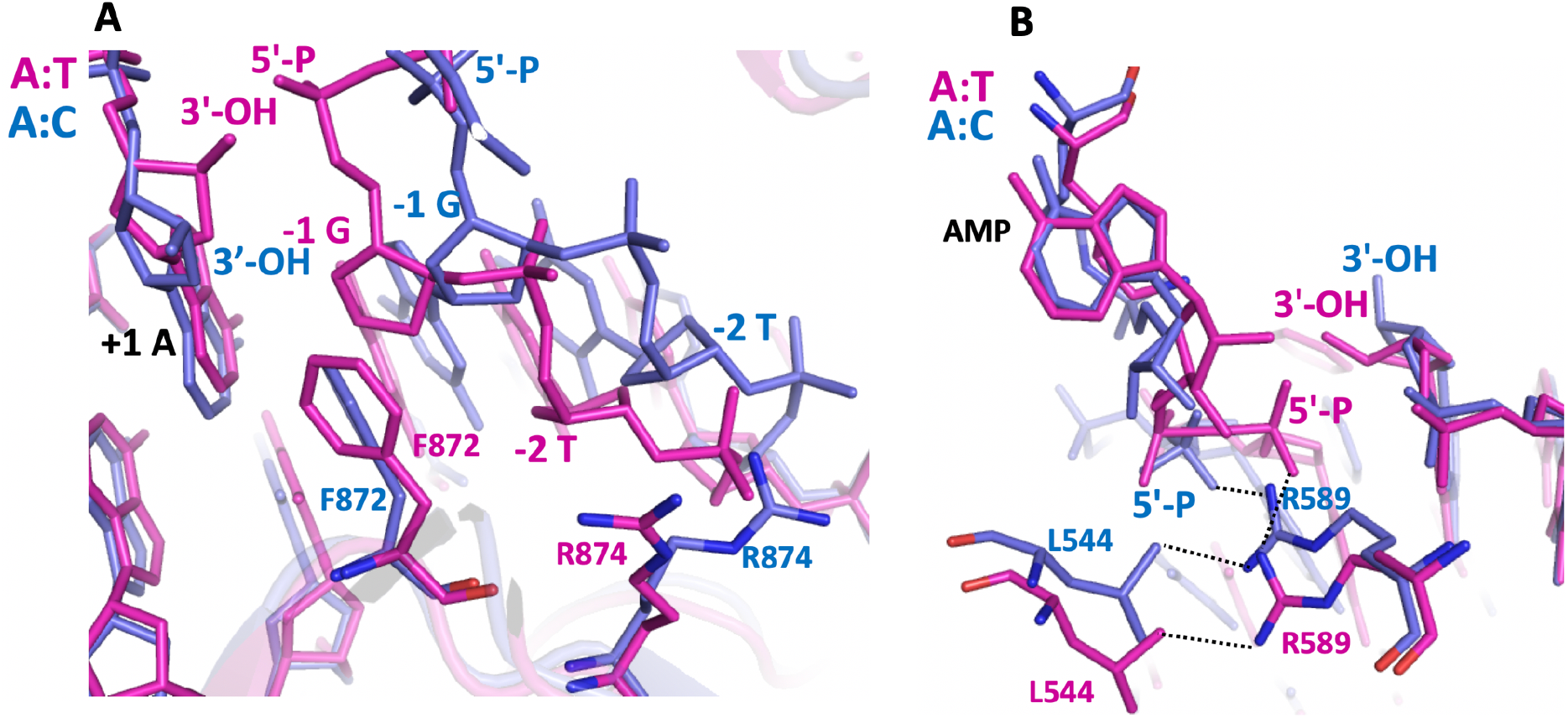
LIG1 active site residues in complex with nick DNA complexes with mismatched ends. Overlay of LIG1 structures bound to the nick DNA duplexes with A:T (magenta) and A:C (blue) show the positions of F872, R874 (**A**) and R589, L544 (**B**). F872 and the neighboring R874 make direct DNA contacts with nucleotides -1G and -2T, respectively. R589 is positioned close to 5′-PO_4_ end and makes a contact with L544 side chain.

The previously solved crystal structures of LIG1 revealed Mg^2+^-dependent high-fidelity (Mg^HiFi^) site that is coordinated by the two conserved glutamate residues at the junction between Adenylation [Glu(E)346] and DNA-binding [Glu(E)592] domains of the ligase and in direct interaction with DNA (54). These structures demonstrated that the mutagenesis at Mg^HiFi^ site (E346A/E592A or EE/AA) allows LIG1 to better accommodate a damaged base (8-oxoG) in the active site. In our study, we also used EE/AA mutant for crystallizations and resolving the structures of LIG1/nick with mismatched ends. Therefore, we finally compared our structures with the previously solved structures of LIG1/nick harboring correct (G:C) and damaged (8-oxoG:A) ends at 3′-strand. The overlay of LIG1/G:C with our all three structures (A:T, G:T, and A:C) showed no difference and the presence of Mg^2+^ at high-fidelity side surrounded by E346 and E592 residues (Supplementary Figure 4). The overlay of LIG1/8-oxodG:A with our LIG1/G:T mismatch demonstrated the ligase structure encircling nick DNA and differences in hydrogen bonding characteristic of G:T and 8-oxodG:A base pairs (Supplementary Figure 5).

### Ligation of the nick repair intermediate with 3′-mismatch by LIG1

In order to evaluate the substrate discrimination mechanism of LIG1 against the repair intermediates harboring mismatched ends for which we determined the ligase/nick DNA structures (Figures 1–5), we performed the ligation assays using the nick DNA substrates containing 3′-preinserted mismatches dG:T and dA:C in a reaction mixture including the wild-type or low-fidelity EE/AA mutant of LIG1. In our control reactions, we evaluated the ligation efficiency of nick DNA with 3′-dA:T.

For both LIG1 proteins, at earlier time points of ligation reaction (10-30 s), we observed an efficient ligation of nick DNA substrates with 3′-dA:T (Figure 6A and 6B, lanes 2-5) and 3′-dG:T (Figure 6A and 6B, lanes 7-10) that yielded similar amount of nick sealing products (Figure 6C-D). However, the end joining efficiency of LIG1 is significantly diminished in the presence of dA:C mismatch at the 3′-end of nick DNA (Figure 6A and 6B, lanes 12-15). There was ~90-fold difference in the amount of ligation products between dG:T and dA:C mismatches (Figure 6C). The only difference between wild-type and low-fidelity EE/AA mutant of LIG1 is the formation of DNA intermediates with 5′-adenylate (AMP). We observed more intermediate products accumulated in the presence of 3′-dA:C and 3′-dG:T mismatches by LIG1 EE/AA and wild-type, respectively (Supplementary Figure 6). However, even for longer time points of ligation reaction incubation (10 min), we observed very efficient ligation for the nick DNA substrates containing 3′-dA:T and 3′-dG:T and drastic decrease in nick sealing efficiency of 3′-dA:C by both wild-type and EE//AA mutant of LIG1 (Supplementary Figures 7 and 8).

**Figure 6.**
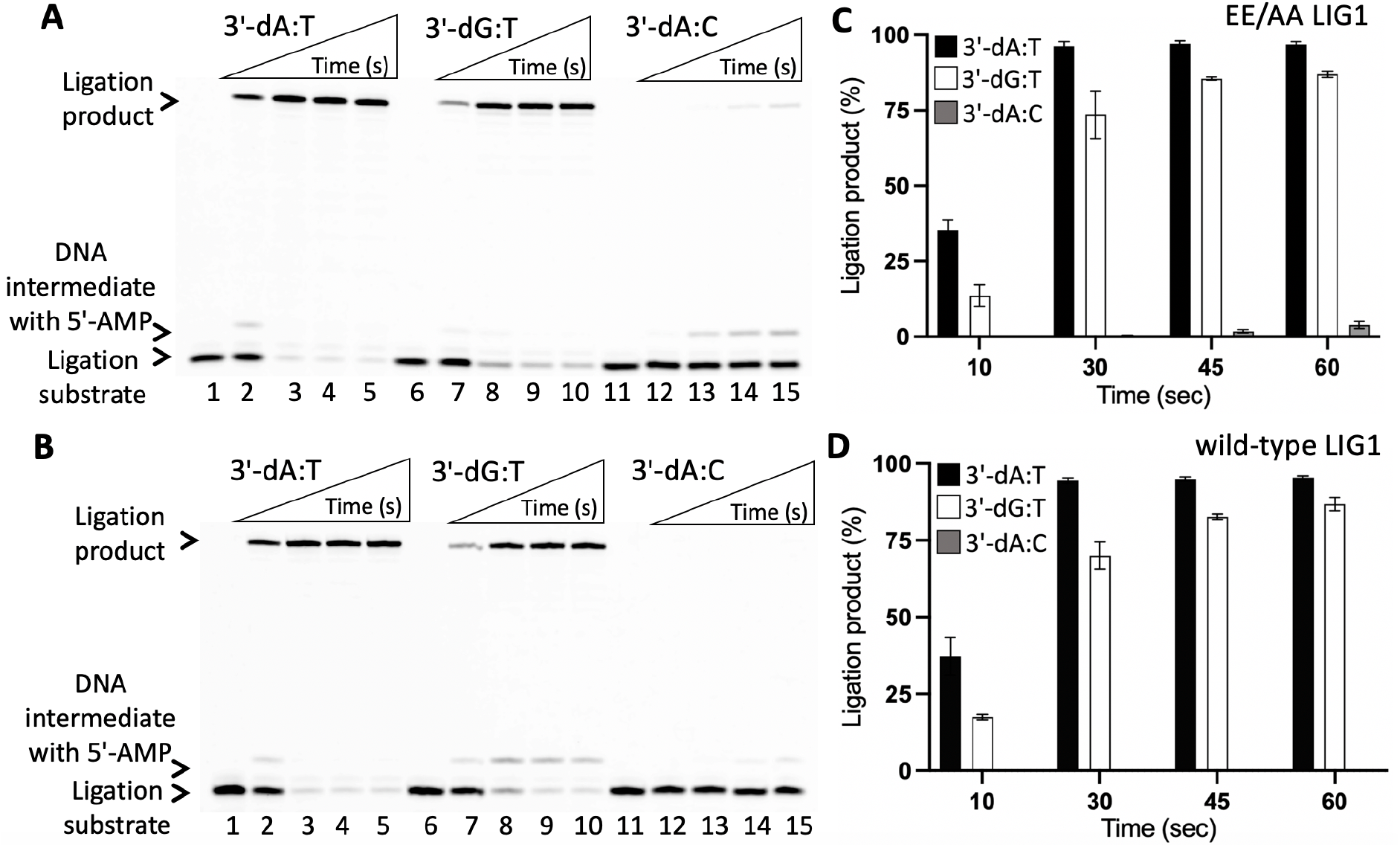
Ligation of nick repair intermediate with G:T and A:C mismatches by LIG1. (**A,B**) Lanes 1, 6, and 11 are the negative enzyme controls of the nick DNA substrates with 3′-preinserted dA:T, dG:T, and dA:C, respectively. Lanes 2-5, 7-10, and 12-15 are the reaction products for nick sealing of DNA substrates with 3′-preinserted dA:T, dG:T, and dA:C, respectively, by EE/AA (A) and wild-type (B) of LIG1, and correspond to time points of 10, 30, 45, and 60 sec. (**C,D**) The graphs show time-dependent changes in the amount of ligation products. The data represent the average of three independent experiments ± SD.

In our previous studies, we reported that polβ 8-oxodGTP insertion leads to ligase failure and the mutation (EE/AA) at the high-fidelity site of LIG1 results in the mutagenic ligation of nick repair intermediate with an inserted 8-oxodGMP (35). In the present study, we also compared the DNA end-joining efficiency of LIG1 against the repair intermediates with 3′-preinserted mismatch and damaged base that mimic DNA polymerase mismatch (dGTP:C) and oxidized nucleotide (8-oxodGTP:A) insertion products, respectively. For this purpose, we used the nick DNA substrates with 3′-preinserted dG:T and 8-oxodG:A in the ligation reaction as described above. In consistent with our previous studies (32,37–40), we found that LIG1 EE/AA can ligate the nick DNA substrates with 3′-dG:T and 3′-8-oxodG:A efficiently (Supplementary Figure 9A, lanes 2-5 and 7-10, respectively). We did not obtain significant difference in the amount of ligation products (Supplementary Figure 9B). Similarly, wild-type LIG1 shows similar end-joining efficiency for both nick DNA substrates containing mismatched and damaged bases at 3′-end (Supplementary Figure 9C, lanes 2-5 and 7-10, respectively) with time-dependent increase in the amount of ligation products (Supplementary Figure 9D). We also observed the accumulation of DNA intermediates with 5′-AMP (Supplementary Figure 10).

### Ligation of polβ mismatch nucleotide insertion products by LIG1

The discrimination of the repair intermediates by DNA ligases can impair progression of BER pathway when mismatch nucleotides are inserted by polβ (26–36). In order to further understand the effect of mismatches on the interplay between polβ and LIG1 at the downstream steps of the BER pathway, we investigated the ligation of polβ correct (dGTP:C) versus mismatch (dGTP:T and dATP:C) nucleotide insertion products in a coupled reaction including polβ and LIG1 (wild-type or EE/AA mutant). For this purpose, we used one nucleotide gap DNA substrates with template C or T.

Consistent with our previous findings (32), we observed that the repair products after polβ dGTP:T insertion were ligated efficiently by wild-type LIG1 (Figure 7A, lanes 7-10). These products are similar to the end joining products of nick repair intermediates after polβ correct dGTP:C insertion (Figure 7A, lanes 2-5). The amount of ligation products in a coupled reaction after polβ dGTP:T insertion was relatively lower when compared with the ligation products after polβ dGTP:C insertion (Figure 7B). However, in the coupled reaction where we tested the ligation of polβ dATP mismatch insertion opposite C by wild-type LIG1 (Figure 7A, lanes 12-15), there was no ligation products (Figure 7B).

**Figure 7.**
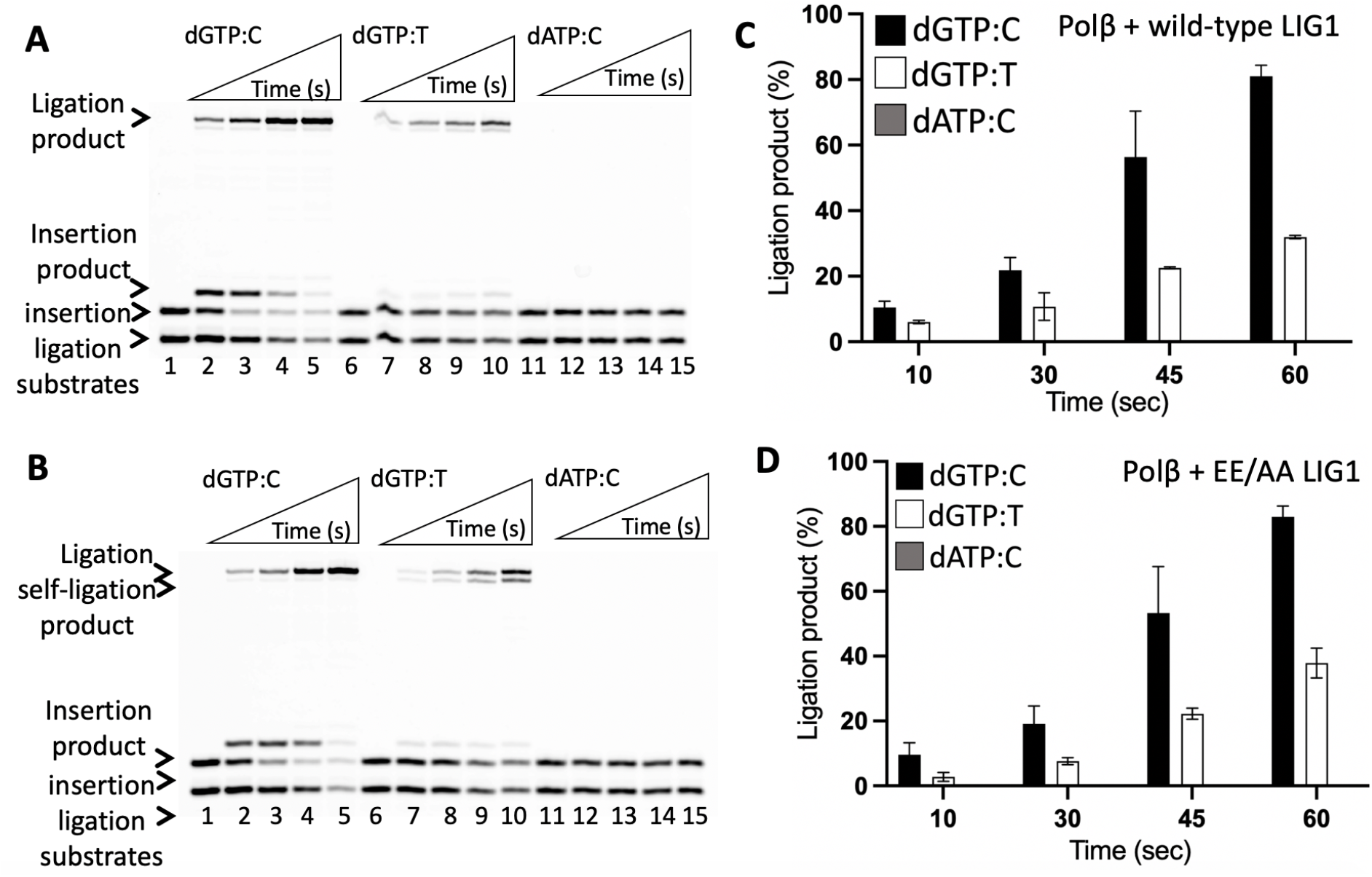
Ligation of polβ mismatch nucleotide insertion products by LIG1. (**A,B**) Lanes 1, 6, and 11 are the negative enzyme controls of the one nucleotide gap DNA substrate with template C, T, and C, respectively. Lanes 2-5, 7-10, and 12-15 are the reaction products for the ligation of polβ dGTP:C, dGTP:T, dATP:C insertions by wild-type (A) and EE/AA (B) of LIG1, respectively, and correspond to time points of 10, 30, 45, and 60 sec. (**C,D**) The graphs show time-dependent changes in the amount of ligation products. The data represent the average of three independent experiments ± SD.

For low-fidelity LIG1 EE/AA mutant, we also obtained efficient end joining of polβ dGTP:T insertion product (Figure 7C, lanes 7-10). This was similar to the ligation products of polβ correct dGTP insertion opposite C in the control reactions (Figure 7C, lanes 2-5). Interestingly, the products of self-ligation (*i.e*., end joining of one nucleotide gap DNA itself) were appeared simultaneously with a complete ligation of nicked insertion products (Figure 7C, compare lines 5 and 10). Similar to the wild-type enzyme, LIG1 EE/AA mutant was not able to ligate the repair intermediate after polβ dATP:C mismatch insertion (Figure 7C, lanes 12-15). The amount of ligation products after polβ insertions for dGTP:C was higher than those after dGTP:T insertion (Figure 7D).

### Interplay between APE1 and LIG1 during processing of the nick repair intermediates with mismatched ends

DNA repair intermediates with a damaged or mismatched base at 3′-end could block the pathway coordination and become persistent DNA strand-breaks if not repaired (14). In our previous studies, we reported the role of DNA end-processing proteins Aprataxin (APTX) and Flap Endonuclease 1 (FEN1) in cleaning an adenylate (AMP) block from 5′-end of the ligation failure products (23–25). In order to further investigate the processing of mutagenic nick repair products with an inserted mismatched bases at 3′-end, we examined the role of APE1 as a compensatory DNA end-processing enzyme (69). For this purpose, we evaluated the 3′-5′ exonuclease activity of APE1 in a reaction mixture that includes the nick DNA substrates with 3′-preinserted dG:T and dA:C mismatches. We did not observe a significant difference in the mismatch base removal efficiency of APE1 between 3′-dG:T and 3′-dA:C mismatches (Figure 8A). Our results demonstrated that APE1 can remove 3′-dG and 3′-dA bases from the nick DNA substrates with template base T and C, respectively (Supplementary Figure 11).

**Figure 8.**
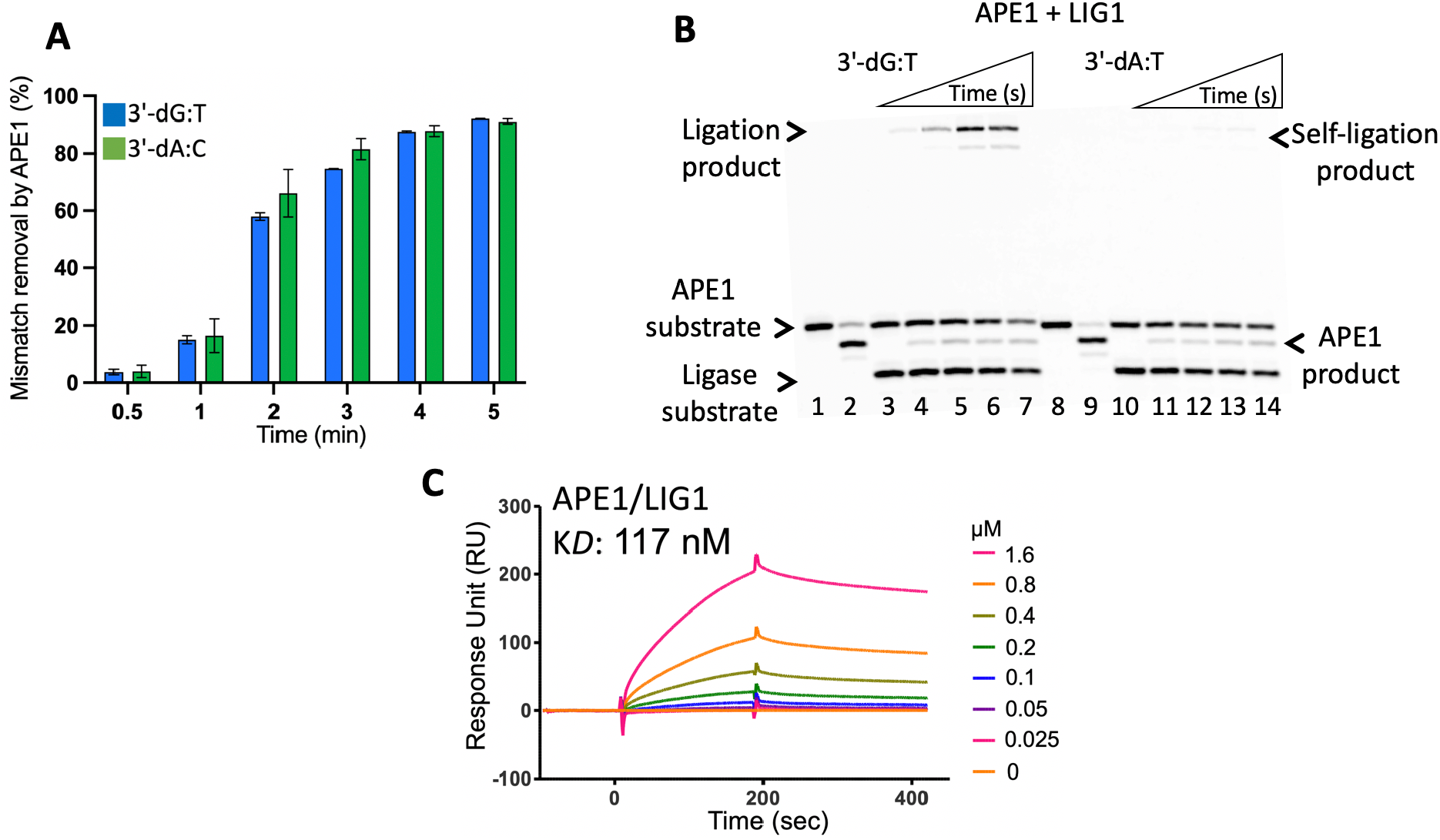
APE1 and LIG1 interplay on the repair intermediate with G:T and A:C mismatches. (**A**) APE1 activity on the removal of a mismatched bases from the nick repair intermediates with 3′-preinserted dG:T and dA:C. The graph shows time-dependent changes in the amount of APE1 excision products. The data represent the average of three independent experiments ± SD. **(B**) APE1 mismatch removal coupled to ligation by LIG1. Lanes 1 and 8 are the negative enzyme controls and lanes 2 and 9 are APE1 mismatch removal products for the nick DNA substrates with 3′-preinserted dG:T and dA:C, respectively. Lanes 3-7 and 10-14 are the reaction products for APE1 mismatch removal and nick sealing of DNA substrates with 3′-preinserted dG:T and dA:C, respectively, and correspond to time points of 10, 30, 45, 60, and 120 sec. **(C**) Real-time protein-protein interaction analysis between APE1 and LIG1.

In line with our observations that demonstrate the end joining ability of LIG1 on the mismatch-containing nick repair intermediates (Figure 6), we further investigated the processing of the nick DNA substrates with 3′-dG:T and 3′-dA:C mismatches in coupled reactions including both APE1 and LIG1 to test the efficiency of mismatch removal and ligation simultaneously. Our results demonstrated that APE1 mismatch removal products were accumulated along with the ligation products for the nick DNA substrates with 3′-dG:T mismatch (Figure 8B, lanes 2-5). However, we mainly observed the products of 3′-dA mismatch removal by APE1 from the nicked DNA substrate with 3′-dA:C (Figure 8B, lanes 7-10).

Lastly, we quantitatively monitored the real time kinetics of protein-protein interaction between APE1 and LIG1 by Surface Plasmon Resonance (SPR) assay where the interacting protein partner of APE1 was immobilized on CM5 biosensors onto which LIG1 protein was respectively passed as an analyte. Our results, for the first time, showed protein-protein interaction with the equilibrium binding constant (K*D*: 117 nM) between APE1 and LIG1 (Figure 8C). In previously published studies, the physical interactions for APE1 were reported for the core BER proteins such as DNA glycosylase MYH, polβ, and XRCC1 (19,20). Thermodynamics and domain mapping studies also showed that polβ interacts with the N-terminal noncatalytic part of LIG1 (70,71). Overall, it seems that the interplay within the multi-protein BER process when bound to a repair intermediate with an incompatible end could affect the repair efficiency at the downstream steps.

## Discussion

The BER is a critical process for preventing the mutagenic and lethal consequences of complex types of lesions (15). The fidelity of BER requires tightly coordinated series of enzymatic steps, which is critical to prevent the release and accumulation of toxic and mutagenic single-strand break intermediates that contribute to genome instability (16–18). Although structural and biochemical studies have provided extensive evidence for sequential substrate-product shuttling for faithful BER coordination at earlier steps of the repair pathway, the molecular coordination at the downstream steps involving polβ gap filling and final DNA ligation by ligase I or IIIα remains largely unknown.

Kinetic, structural, and computational studies have demonstrated that the polβ active site undergoes a conformational change to form the precatalytic closed ternary complex in the presence of a Watson-Crick base pairing, known as an induced-fit mechanism, exhibits diverse mismatch-induced conformational distortions that is dependent on the architecture of the mismatched template primer (58–68). The largest distortion has been reported for A:C mismatch where O3′ of the primer terminus sugar is positioned away from the active site by precluding direct template base interactions, which results in a loss of one hydrogen bond, and therefore, effectively deters further DNA synthesis (64,65). Polβ structures also revealed that the polymerase active site with an inserted dGMP opposite T escapes mismatch discrimination through ionization of the wobble base pair and exhibits a Watson-Crick-like conformation in a closed state (58–68). Despite of these polβ binary and ternary complex structures with a variety of mismatches, how downstream proteins, particularly DNA ligase, engage with the mismatch-containing repair intermediates is still lacking. In our present study, we determined at atomic resolution the features of the DNA substrate and LIG1 interaction that dictate accurate versus mutagenic outcomes to define the critical elements of accurate BER at final steps of the repair pathway.

In this work, we used structural and biochemical approaches to elucidate the mechanisms by which LIG1 discriminates against mutagenic 3′-mismatches that could be formed during prior polβ mismatch nucleotide insertion step of the BER pathway. Our study reveals that LIG1 discriminates 3′ termini depending on the architecture of mismatched ends at during steps 1 (ligase adenylation) and step 2 (AMP transfer to DNA) of the ligation reaction. The structure of LIG1 bound to nick DNA duplexes with A:T and G:T showed that the ligase can ligate G:T mismatch with two hydrogen bonds and a base-pair size that is nearly indistinguishable from that of a Watson-Crick A:T base pair. Both structures refer to LIG1 bound to AMP-DNA nick as step 2 product of ligase reaction. We also showed efficient nick sealing of DNA substrates with 3′-preinserted dA:T and dG:T to a significant degree with high efficiency in ligation reactions *in vitro*. Importantly, our structures revealed that the LIG1 active site is quite rigid and ligation differences between nick DNA substrates with G:T and A:C mismatches likely result from altered DNA conformations. Crystal structure of LIG1/nick DNA with A:C mismatch demonstrated that the mismatched termini distorts the conformations of 3′- and 5′-strands of a nick. Overall, our results demonstrate that the extent to which G:T versus A:C mismatch discrimination by LIG1 counteracts the polymerase-promoted mutagenesis products distinctly at the downstream steps of BER pathway.

All mammalian and bacterial ATP-dependent DNA ligases contain a highly conserved the catalytic core consisting of the oligonucleotide binding domain (OBD) and the adenylation (AdD) or nucleotidyl transferase (NTase) domain (3–5). Despite of this C-terminal core architecture, they display different fidelity profiles depending on the type (human ligase I, IIIα, or IV) and source of DNA ligase (54–57,73–95). Metal ions play important roles in steps 1 and 3 of the ligation reaction (1–13). The catalytic Mg^2+^ ion deprotonates the lysine nucleophile to activate it to attack on the ATP α-phosphate and stabilize the pentavalent transition state formed during step 1, while the metal ion activates 3′-OH nucleophilic attack on the 5′-phosphate and form the pentavalent transition during final nick sealing step 3 (73–95). We crystallized LIG1 EE/AA in complex with matched A:T and mismatched G:T and A:C nick DNA under similar conditions in the absence of metal ion. Most structures of ATP-dependent DNA ligases have no divalent cation in the refined model and the mechanism of step 1 during which LIG1 engages with catalytic Mg^2+^ and ATP remains unclear at atomic resolution. Previously solved crystal structures of DNA ligase/ATP and ligase/ATP/ Mg^2+^ complexes for ATP-dependent ligases from other sources such as *Chlorella virus* DNA ligase, T4 DNA ligase, *Mycobacterium tuberculosis* LigD, and *Pyrococcus furiosus* DNA ligase in a noncovalent complex with AMP enlighten the requirement of metal ion for the ligase adenylation (83–94). The present study represents a human LIG1 bound to ATP (LIG1-AMP) in step 1 of ligation reaction while engaging with nick DNA harboring A:C mismatched termini at 3′-strand.

Based on our structural and biochemical results, according to our model (Supplementary Scheme 3), we hypothesized that the LIG1-nick repair intermediate with a poor mismatch (A:C) versus a good match (A:T) architecture could serve as a structural fidelity checkpoint at which the efficiency of repair pathway coordination is mediated at the final ligation step. Furthermore, the polβ mismatched versus matched-substrate/product complex could also determine the fate of substrate-product channeling that can deter or favor final nick sealing step in the BER pathway. This phenomenon could provide an opportunity for 3′-proofreading by APE1. It has been also reported in the structure studies that the nick with a mismatched base exhibits distinct features where 3′-end slides into the APE1 active site and a mismatched end is stabilized by protein contacts (96). It is also likely that other 3′-5′ exonucleases can also provide a proofreading function at the downstream steps of BER pathway for 3′-end cleaning (97). Importantly, our study uncovered a mutagenic event where the architecture of LIG1/nick duplex with G:T mismatch in the active site forms a premutagenic structural intermediate. In our previous study, we reported that DNA polymerase (pol) μ dGTP mismatch insertion opposite T during gap filling repair synthesis in the nonhomologous end joining (NHEJ) pathway is effectively coupled with ligation by DNA ligase IV, resulting in the formation of promutagenic NHEJ intermediate (30). Recently, the crystal structures of pol μ misincorporating dGTP on gap DNA substrate containing template T revealed its highly mutagenic base pairing role at the 3′ end of the gap during NHEJ (98). Further structure/function studies with both BER DNA ligases (ligase I and IIIα) are required for all other possible non-canonical base pairings at 3′-end of a nick to comprehensively understand the ligase strategies against the mutagenic repair intermediates that could be formed due to aberrant BER function of polβ such as cancer-associated variants with slower or lack of gap filling activity and reduced fidelity (37–46).

## Methods

### Preparation of DNA substrates for crystallization and BER assays

Oligodeoxyribonucleotides with and without a 6-carboxyfluorescein (FAM) label were obtained from Integrated DNA Technologies (IDT). DNA substrates were prepared as described previously (23–36). The nick DNA substrates containing 3′-preinserted correct (dA:T), mismatch (dG:T and dA:C), or damaged (8-oxodG:A) ends with a FAM label at 3′-end were used for DNA ligation assays in a reaction mixture including LIG1 (wild-type or EE/AA mutant) alone (Supplementary Table 1). The one nucleotide gap DNA substrates containing FAM labels at both 3′-and 5′-ends were used for the coupled assays to observe the ligation of polβ correct or mismatch nucleotide insertion products by LIG1 (wild-type or EE/AA mutant) in the same reaction mixture including both polβ and LIG1 (Supplementary Table 2). The nick DNA substrates containing 3′-preinserted correct (dA:T) or mismatch (dG:T and dA:C) with a FAM label at 5′-end were used for APE1 exonuclease assays in a reaction mixture including APE1 alone (Supplementary Table 3). The nick mismatch containing DNA substrates with FAM labels at both 3′- and 5′-ends were used in the coupled assays to observe APE1 mismatch removal and ligation in the same reaction mixture including both APE1 and LIG1 (Supplementary Table 4). For LIG1 X-ray crystallography studies, the nick DNA substrates containing correct A:T and mismatch G:T and A:C ends were prepared by annealing upstream, downstream, and template primers (Supplementary Table 5).

### Protein purifications

Human his-tag recombinant full-length (1-918) wild-type DNA ligase I (LIG1) and C-terminal (△261) E346A/E592A (EE/AA) mutant were purified as described previously (23–36). Briefly, the proteins were overexpressed in Rosetta (DE3) pLysS *E. coli* cells (Millipore Sigma) and grown in Terrific Broth (TB) media with kanamycin (50 μgml^−1^) and chloramphenicol (34 μgml^−1^) at 37 °C. Once the OD was reached to 1.0, the cells were induced with 0.5 mM isopropyl β-D-thiogalactoside (IPTG) and the overexpression was continued for overnight at 28 °C. After the centrifugation, the cell was lysed in the lysis buffer containing 50 mM Tris-HCl (pH 7.0), 500 mM NaCl, 20 mM imidazole, 10% glycerol, 1 mM PMSF, an EDTA-free protease inhibitor cocktail tablet by sonication at 4 °C. The lysate was pelleted at 16,000 x rpm for 1h at 4 °C. The cell lysis solution was filter clarified and then loaded onto a HisTrap HP column (GE Health Sciences) that was previously equilibrated with the binding buffer including 50 mM Tris-HCl (pH 7.0), 500 mM NaCl, 20 mM imidazole, 10% glycerol. The column was washed with the binding buffer and then followed by washing buffer containing 50 mM Tris-HCl (pH 7.0), 500 mM NaCl, 35 mM imidazole, 10% glycerol. The protein was finally eluted with an increasing imidazole gradient 0-500 mM at 4 °C. The collected fractions were then subsequently loaded onto HiTrap Heparin (GE Health Sciences) column that was equilibrated with binding buffer containing 50 mM Tris-HCl (pH 7.0), 50 mM NaCl, 0.2 mM EDTA, and 10% glycerol, and protein is eluted with a linear gradient of NaCl up to 1 M. The LIG1 protein was further purified by Resource Q and finally by Superdex 200 10/300 (GE Health Sciences) columns in the buffer containing 20 mM Tris-HCl (pH 7.0), 200 mM NaCl, and 5% glycerol.

Human wild-type AP-Endonuclease 1 (APE1) with his-tag (pET-24b) was overexpressed and purified as previously described (23–36). Briefly, the protein was overexpressed in BL21(DE3)*E.coli* cells (Invitrogen) in Lysogeny Broth (LB) media at 37 °C for 8 h, induced with 0.5 mM IPTG and the overexpression was continued for overnight at 28 °C. After the cells were harvested, lysed at 4 °C, and then clarified as described above. The supernatant was loaded onto a HisTrap HP column (GE Health Sciences) and purified with an increasing imidazole gradient (0-300 mM) elution at 4 °C. The collected fractions were then subsequently loaded onto a HiTrap Heparin column (GE Health Sciences) with a linear gradient of NaCl up to 1 M. The recombinant APE1 were then further purified by Superdex 200 increase 10/300 chromatography (GE Healthcare) in the buffer containing 20 mM Tris-HCl (pH 7.0), 200 mM NaCl, and 1 mM DTT.

Human wild-type polβ with GST-tag (pGEX-6p-1) were overexpressed and purified as previously described (23–36). Briefly, the protein was overexpressed in One Shot BL21(DE3)pLysS *E.coli* cells (Invitrogen) in LB media at 37 °C for 8 h, induced with 0.5 mM IPTG, and the overexpression was continued for overnight at 28 °C. The cells were then grown overnight at 16 °C. After cell lysis at 4 °C by sonication in the lysis buffer containing 1X PBS (pH 7.3), 200 mM NaCl, and 1 mM DTT, and cOmplete Protease Inhibitor Cocktail (Roche), the lysate was pelleted at 16,000 x rpm for 1 h and then clarified by centrifugation and filtration. The supernatant was loaded onto a GSTrap HP column (GE Health Sciences) and purified with the elution buffer containing 50 mM Tris-HCl (pH 8.0) and 10 mM reduced glutathione. In order to cleave a GST-tag, the recombinant protein was incubated with PreScission Protease (GE Health Sciences) for 16 h at 4 °C in the buffer containing 1X PBS (pH 7.3), 200 mM NaCl, and 1 mM DTT. After the cleavage, the polβ protein was subsequently passed through a GSTrap HP column, and the protein without GST-tag were then further purified by loading onto Superdex 200 gel filtration column (GE Health Sciences) in the buffer containing 50 mM Tris-HCl (pH 7.5) and 400 mM NaCl. All proteins purified in this study were dialyzed against storage buffer including 25 mM Tris-HCl (pH 7.0), 200 mM NaCl, concentrated, frozen in liquid nitrogen, and stored at -80 °C. Protein quality was evaluated onto 10% SDS-PAGE, and the protein concentration was measured using absorbance at 280 nm.

### Crystallization and structure determination

LIG1 C-terminal (△261) EE/AA mutant was used for crystals production. All the LIG1-DNA complex crystals were grown at 20 °C using the hanging drop method. LIG1 (at 26 mgml^−1^ LIG1)/DNA complex solution was prepared in 20 mM Tris-HCl (pH 7.0), 200 mM NaCl, 1 mM DTT, 0.1 mM EDTA and 1 mM ATP at 1.5:1 DNA:protein molar ratio of nick DNA and then mixed with 1 μl reservoir solution containing 100 mM MES (pH 6.6), 100 mM lithium acetate, and 20% (w/v) polyethylene glycol PEG3350. All crystals grew in 1-2 days and they were washed in the reservoir solution with 20% glycerol and flash cooled in liquid nitrogen for data collection. The crystals were maintained at 100 K during X-ray diffraction data collection using the beamline 7B2 at Cornell High Energy Synchrotron Source (CHESS). The diffraction images were indexed and integrated using HKL2000. All structures were solved by the molecular replacement method using PHASER using PDB entry 6P0D as a search model (54). Iterative rounds of model building in COOT and refinement with PHENIX or REFMAC5 were used to produce the final models (99–103). All structural images were drawn using PyMOL (The PyMOL Molecular Graphics System, V0.99, Schrödinger, LLC). Detailed crystallographic statistics are provided in Table 1.

### DNA ligation assays

The ligation assays using the nick DNA substrates containing 3′-preinserted correct dA:T, mismatched dG:T or dA:C, and damaged 8-oxodG:A ends were performed (Supplementary Scheme 1A) as described previously (23–36). Briefly, the ligation assays were performed in a mixture containing 50 mM Tris-HCl (pH 7.5), 100 mM KCl, 10 mM MgCl_2_, 1 mM ATP, 1 mM DTT, 100 μgml^−1^ BSA, 10% glycerol, and 500 nM DNA substrate in a final volume of 10 μl. The reactions were initiated by the addition of 100 nM LIG1 (wild-type or EE/AA mutant), incubated at 37 °C, and stopped at the time points indicated in the figure legends. The reaction products were then quenched with an equal amount of gel loading buffer containing 95% formamide, 20 mM ethylenediaminetetraacetic acid, 0.02% bromophenol blue and 0.02% xylene cyanol. After incubation at 95 °C for 3 min, the reaction products were separated by electrophoresis on an 18% denaturing polyacrylamide gel. The gels were scanned with a Typhoon PhosphorImager (Amersham Typhoon RGB), and the data were analyzed using ImageQuant software.

### BER assays to measure DNA ligation of polβ nucleotide insertion products

The one nucleotide gap DNA substrates with template A or C were used to test the ligation of polβ nucleotide insertion (correct or mismatch) products in the reaction mixture including polβ and LIG1 (Supplementary Scheme 1B) as described previously (23–36). Briefly, the coupled assays were performed in a mixture containing 50 mM Tris-HCl (pH 7.5), 100 mM KCl, 10 mM MgCl_2_, 1 mM ATP, 1 mM DTT, 100 μgml^−1^ BSA, 10% glycerol, 100 μM dNTP, and 500 nM DNA substrate in a final volume of 10 μl. The reactions were initiated by the addition of pre-incubated enzyme mixture of polβ/LIG1 (100 nM) and incubated at 37 °C for the time points as indicated in the figure legends. The reaction products were then mixed with an equal amount of gel loading buffer, separated by electrophoresis on an 18% denaturing polyacrylamide gel, and analyzed as described above.

### BER assays to measure APE1 exonuclease activity and LIG1 ligation

The nick DNA substrates including 3′-preinserted mismatches dG:T and dA:C were used to examine APE1 proofreading role for removing a mismatched base by its 3′-5′ exonuclease activity (Supplementary Scheme 2A). Briefly, APE1 activity assays were performed in a mixture containing 50 mM HEPES (pH 7.4), 100 mM KCl, 3 mM MgCl_2_, 0.1 mgml^−1^ BSA, and 500 nM DNA substrate in a final volume of 10 μl. The reactions were initiated by the addition of 50 nM APE1, incubated at 37 °C for the time points as indicated in the figure legends, quenched by mixing with 100 mM EDTA, and then mixed with an equal amount of gel loading buffer. The nick DNA substrates including 3′-preinserted mismatches dG:T and dA:C were used for repair assays to test APE1 exonuclease and DNA ligation activities in the same reaction mixture (Supplementary Scheme 2B). Briefly, the repair assays were performed in a mixture containing 50 mM HEPES (pH 7.4), 100 mM KCl, 5 mM MgCl_2_, 1 mM ATP, 0.1 mgml^−1^ BSA, and 500 nM DNA substrate in a final volume of 10 μl. The reactions were initiated by the addition of pre-incubated enzyme mixture including APE1/LIG1 (100 nM), incubated at 37 °C for the time points as indicated in the figure legends. The reaction products were quenched by mixing with 100 mM EDTA and then mixed with an equal amount of gel loading buffer. The reaction products were separated by electrophoresis on an 18% denaturing polyacrylamide gel and analyzed as described above.

### APE1 and LIG1 protein-protein interaction assay

The protein-protein interaction between APE1 and LIG1 was measured by Surface Plasmon Resonance (SPR) in real time using Biacore X-100 (GE Healthcare) as described previously (36). Briefly, one flow cell of the CM5 sensor chip was activated at 25 °C with a 1:1 mixture of 0.2 M EDC and 0.05 M NHS in water, and then APE1 protein was injected over the flow cell in 10 mM sodium acetate at pH 5.0 at a flow rate of 10 μl/min. The binding sites were blocked using 1 M ethanolamine. LIG1 (at the concentration range of 0-1.6 μM) was then injected for 3 min at a flow rate of 30 μl/min in the binding buffer (20 mM HEPES pH 7.4, 150 mM NaCl, 3 mM EDTA and 0.005% (v/v) Surfactant P20). After a dissociation phase for 3-4 min, 0.2% SDS was injected for 30 sec to regenerate the chip surface. Non-specific binding to a blank flow cell was subtracted to obtain corrected sensorgrams. All data were analyzed using BIAevaluation software version 2.0.1 and fitted to a 1:1 (Langmuir) binding model to obtain equilibrium constant (K*D*).

## Supporting information

Supplementary Data for BioRvix 2021-Caglayan

## Data availability

Atomic coordinates and structure factors for the reported crystal structures have been deposited in the RCSB Protein Data Bank under accession numbers 7SUM, 7SXE, 7SX5. All relevant data are available from the authors upon reasonable request.

## Acknowledgements

This work is based upon research conducted at the Center for High Energy X-ray Sciences (CHEXS), which is supported by the National Science Foundation under award DMR-1829070, and the Macromolecular Diffraction at CHESS (MacCHESS) facility, which is supported by award 1-P30-GM124166-01A1 from the National Institute of General Medical Sciences, National Institutes of Health, and by New York State’s Empire State Development Corporation (NYSTAR). The authors thank Jacob E. Combs (McKenna Lab, University of Florida) for his assistance with crystal shipment and data collection.

## Author contributions

Conceptualization M.Ç., methodology and investigation M.Ç., T.Q.; writing-original draft, M.Ç., T.Q.; writing-reviewing and editing, M.Ç., T.Q., R.M.; funding acquisition M.Ç.

## References

1. Timson, D. J., Singleton, M. R. & Wigley, D. B. DNA ligases in the repair and replication of DNA. Mutat. Res. 460, 301–318 (2000).

2. Shuman, S. DNA ligases: progress and prospects. J. Biol. Chem. 284, 17365–17369 (2009).

3. Tomkinson, A. E., Vijayakumar, S., Pascal, J. M. & Ellenberger, T. DNA ligases: structure, reaction mechanism, and function. Chem. Rev. 106, 687–699 (2006).

4. Doherty, A. J. & Suh, S. W. Structural and mechanistic conservation in DNA ligases. Nucleic Acids Res. 28, 4051–4058 (2000).

5. Ellenberger, T. & Tomkinson, A. E. Eukaryotic DNA ligases: Structural and functional insights. Annu. Rev. Biochem. 77, 313–338 (2008).

6. Cherepanov, A. V. & Vries, S. Dynamic mechanism of nick recognition by DNA ligase. Eur. J. Biochem. 269, 5993–5999 (2002).

7. Dickson, K. S., Burns, C. M. & Richardson, J. P. Determination of the free-energy change for repair of a DNA phosphodiester bond. J. Biol. Chem. 275, 15828–15831 (2000).

8. Weiss, B., Thompson, A. & Richardson, C. C. Ezymatic breakage and joining of deoxyribonucleic acid. VII. Properties of the enzyme-adenylate intermediate in the polynucleotide ligase reaction. J. Biol. Chem. 243, 4556–4563 (1968).

9. Singleton, M. R., Hakansson, K., Timson, D. J. & Wigley, D. B. Structure of the adenylation domain of an NAD^+^-dependent DNA ligase. Structure 7, 35–42 (1999).

10. Tomkinson, A. E., Totty, N. F., Ginsburg, M. & Lindahl, T. Location of the active-site for enzyme-adenylate formation in DNA ligases. Proc. Natl. Acad. Sci. USA 88, 400–404 (1991).

11. Yang, S. W. & Chan, J. Y. Analysis of the formation of AMP-DNA intermediate and the successive reaction by human DNA ligases I and II. J. Biol. Chem. 267, 8117–8122 (1992).

12. Taylor, M. R., Conrad, J. A., Wahl, D. & O’Brien, P. J. Kinetic mechanism of human DNA ligase I reveals magnesium-dependent changes in the rate-limiting step that compromise ligation efficiency. J. Biol. Chem. 286, 23054–23062 (2011).

13. Cherepanov, A. V. & Vries, S. Kinetics and thermodynamics of nick sealing by T4 DNA ligase. Eur. J. Biochem. 270, 4315–4325 (2003).

14. Caglayan, M. Interplay between DNA polymerases and DNA ligases: Influence on substrate channeling and the fidelity of DNA ligation. J. Mol. Biol. 431, 2068–2081 (2019).

15. Beard, W. A. et al. Eukaryotic base excision repair: New approaches shine light on mechanism. Ann. Rev. Biochem. 88, 137–162 (2019).

16. Prasad, R. et al. A review of recent experiments on step-to-step “hand-off” of the DNA intermediates in mammalian base excision repair pathways. Mol. Biol. 45, 586–600 (2011).

17. Prasad, R., Shock, D. D., Beard, W. A. & Wilson, S. H. Substrate channeling in mammalian base excision repair pathways: passing the baton. J. Biol. Chem. 285, 40479–40488 (2010).

18. Wilson, S. H. & Kunkel, T. A. Passing the baton in base excision repair. Nat. Struct. Biol. 7, 176–178 (2000).

19. Moor, N. A. et al. Quantitative characterization of protein-protein complexes involved in base excision DNA repair. Nucleic Acids Res. 43, 6009–6022 (2015).

20. Moor, N. A. & Lavrik, O. I. Protein-protein interactions in DNA base excision repair. Biochemistry 83, 411–422 (2018).

21. Caglayan, M. & Wilson, S. H. Oxidant and environmental toxicant-induced effects compromise DNA ligation during base excision DNA repair. DNA Repair 35, 85–89 (2015).

22. Beard, W. A. et al. Efficiency of correct nucleotide insertion governs DNA polymerase fidelity. J. Biol. Chem. 277, 47393–47398 (2002).

23. Caglayan, M. et al. Role of polymerase β in complementing aprataxin deficiency during abasic-site base excision repair. Nat. Struct. Mol. Biol. 21, 497–499 (2014).

24. Caglayan, M. et al. Complementation of aprataxin deficiency by base excision repair enzymes. Nucleic Acids Res. 43, 2271–2281 (2015).

25. Caglayan, M. et al. Complementation of aprataxin deficiency by base excision repair enzymes in mitochondrial extracts. Nucleic Acids Res. 45, 10079–10088 (2017).

26. Caglayan, M. et al. Oxidized nucleotide insertion by pol β confounds ligation during base excision repair. Nat. Commun. 8, 14045 (2017).

27. Caglayan, M. & Wilson, S. H. Role of DNA polymerase β oxidized nucleotide insertion in DNA ligation failure. J. Radiat. Res. 58, 603–607 (2017).

28. Horton, J. K. et al. XRCC1 phosphorylation affects aprataxin recruitment and DNA deadenylation activity. DNA Repair 64, 26–33 (2018).

29. Prasad, R. et al. DNA polymerase β: The missing link of the base excision repair machinery in mammalian mitochondria. DNA Repair 60, 77–88 (2017).

30. Caglayan, M. & Wilson, S. H. Pol μ dGTP mismatch insertion opposite T coupled with ligation reveals a promutagenic DNA intermediate during double strand break repair. Nat. Comm. 9, 4213 (2018).

31. Caglayan M. Pol μ ribonucleotide insertion opposite 8-oxodG facilitates the ligation of premutagenic DNA repair intermediate. Sci. Rep. 10, 940 (2020).

32. Caglayan, M. The ligation of pol β mismatch insertion products governs the formation of promutagenic base excision DNA repair intermediates. Nucleic Acids Res. 48, 3708–3721 (2020).

33. Caglayan, M. Pol β gap filling, DNA ligation and substrate-product channeling during base excision repair opposite oxidized 5-methylcytosine modifications. DNA Repair 95,102945 (2020).

34. Tang, Q., Kamble, P. & Caglayan, M. DNA ligase I variants fail in the ligation of mutagenic repair intermediates with mismatches and oxidative DNA damage. Mutagenesis 35, 391–404 (2020).

35. Kamble, P., Hall, K., Chandak, M., Tang, Q. & Caglayan, M. DNA ligase I fidelity the mutagenic ligation of pol β oxidized and mismatch nucleotide insertion products in base excision repair. J. Biol. Chem. 296,100427 (2021).

36. Tang, Q. & Caglayan, M. The scaffold protein XRCC1 stabilizes the formation of polβ/gap DNA and ligase IIIα/nick DNA complexes in base excision repair. J. Biol. Chem. 297, 101025 (2021).

37. Donigan, K. A. et al. Human polymerase β is mutated in high percentage of colorectal tumors. J. Biol. Chem. 287, 23830–23839 (2012).

38. Starcevic, D. et al. Is there a link between DNA polymerase β and cancer? Cell Cyle 3, 998–1001 (2004).

39. Alnajjar, K. S. et al. A change in the rate-determining step of polymerization by the K289M DNA polymerase β cancer-associated variant. Biochemistry 56, 2096–2105 (2017).

40. Shah, A. M. et al. Variants of DNA polymerase β extend mispaired DNA due to increased affinity for nucleotide substrate. Biochemistry 42, 10709–10717 (2003).

41. Sweasy, J. B. et al. Expression of DNA polymerase β cancer-associated variants in mouse cells results in cellular transformation. Proc. Natl. Acad. Sci. U S A 102, 14350–14355 (2005).

42. Donigan, K. A. et al. The human gastric cancer-associated DNA polymerase beta variant D160N is a mutator that indices cellular transformation. DNA Repair 12, 381–390 (2012).

43. Lang, T. et al. The E295K DNA polymerase β gastric cancer-associated variant interferes with base excision repair and induces cellular transformation. Mol. Cell Biol. 27, 5587–5596 (2007).

44. Donigan, K. A. et al. DNA polymerase β variant Ile260Met generates global gene expression changes related to cellular transformation. Mutagenesis 27, 683–691 (2012).

45. Yamtich, J. et al. A germline polymorphism of DNA polymerase β induces genomic instability and cellular transformation. PLOS Gen. 8, e10003052 (2012).

46. Nemec, A. A. et al. The S229L colon tumor-associated variant of DNA polymerase β induces cellular transformation as a result of decreased polymerization efficiency. J. Biol. Chem. 289, 13708–13716 (2014).

47. Burgers, P. M. J. & Kunkel, T. A. Eukaryotic DNA replication fork. Annu. Rev. Biochem. 86, 417–438 (2017).

48. Modrich, P. DNA mismatch correction. Annu. Rev. Biochem. 56, 435–466 (1987).

49. Topal, M. D. & Fresco, J. R. Complementary base pairing and the origin of substitution mutations. Nature 263, 285–289 (1976).

50. Bebenek, K., Pedersen, L. C. & Kunkel, T. A. Replication infidelity via a mismatch with Watson-Crick geometry. Proc. Natl. Acad Sci. U S A 108, 1862–1867 (2011).

51. Wang, W., Hellinga, H. W. & Beese, L. S. Structural evidence for the rare tautomer hypothesis of spontaneous mutagenesis. Proc. Natl. Acad Sci. U S A 108, 17644–17648 (2011).

52. Harris, V. H. et al. The effect of tautomeric constant on the specificity of nucleotide incorporation during DNA replication: support for the rare tautomer hypothesis of substitution mutagenesis. J. Mol. Biol. 326, 1389–1401 (2003).

53. Kimsey, I. J. et al. Visualizing transient Watson-Crick-like mispairs in DNA and RNA duplexes. Nature 519, 315–320 (2015).

54. Tumbale, P. P. et al. Two-tiered enforcement of high-fidelity DNA ligation. Nat. Commun. 10, 5431 (2019).

55. Pascal, J. M. et al. Human DNA ligase I completely encircles and partially unwinds nicked DNA. Nature 432, 473–478 (2004).

56. Williams, J. S. et al. High-fidelity DNA ligation enforces accurate Okazaki fragment maturation during DNA replication. Nat. Commun. 12, 482 (2021).

57. Jurkiw, T. J. et al. LIG1 syndrome mutations remodel a cooperative network of ligand binding interactions to compromise ligation efficiency. Nucleic Acids Res. 49, 1619–1630 (2021).

58. Beard, W. A. & Wilson, S. H. Structural insights into the origins of DNA polymerase fidelity. Structure 11, 489–496 (2003).

59. Batra, V. K., Beard, W. A., Shock, D. D., Pedersen, L. C. & Wilson, S. H. Nucleotide-induced DNA polymerase active site motions accommodating a mutagenic DNA intermediate. Structure 13, 1225–1233 (2005).

60. Beard, W. A. et al Enzyme-DNA interactions required for efficient nucleotide incorporation and discrimination in human DNA polymerase β. J. Biol. Chem., 271, 12141–12144 (1996).

61. Johnson, K. A. Role of induced fit in enzyme specificity: a molecular forward/reverse switch. J. Biol. Chem., 283, 26297–26301 (2008).

62. Sawaya, M. R., Prasad, R., Wilson, S. H., Kraut, J. & Pelletier, H. Crystal structures of human DNA polymerase beta complexed with gapped and nicked DNA: evidence for an induced fit mechanism. Biochemistry 36, 11205–11215 (1997).

63. Krahn, J. M., Beard, W. A. & Wilson, S. H. Structural insights into DNA polymerase β deterrents for misincorporation support an induced-fit mechanism for fidelity. Structure 12, 1823–1832 (2004).

64. Batra, V. K., Beard, W. A., Pedersen, L. C. & Wilson, S. H. Structures of DNA polymerase mispaired DNA termini transitioning to pre-catalytic complexes support an induced-fit fidelity mechanism. Structure 24, 1863–1875 (2016).

65. Batra, V. K., Beard, W. A., Shock, D. D., Pedersen, L. C. & Wilson, S. H. Structures of DNA polymerase β with active-site mismatches suggest a transient abasic site intermediate during misincorporation. Mol. Cell. 30, 315–324 (2008).

66. Beard, W. A., Shock, D. D., Yang, X. P., DeLauder, S. F. & Wilson, S. H. Loss of DNA polymerase β stacking interactions with templating purines, but not pyrimidines, alters catalytic efficiency and fidelity. J. Biol. Chem. 277, 8235–8242 (2002).

67. Koag, M. C., Nam, K. & Lee, S. The spontaneous replication error and the mismatch discrimination mechanisms of human DNA polymerase β. Nucleic Acids Res. 42, 11233–11245 (2014).

68. Ahn, J., Kraynov, V. S., Zhong, X., Werneburg, B. G. & Tsai, M. D. DNA polymerase β: effects of gapped DNA substrates on dNTP specificity, fidelity, processivity and conformational changes. Biochem. J. 331, 79–87 (1998).

69. Whitaker, A. M. & Freudenthal, B. D. APE1: A skilled nucleic acid surgeon. DNA Repair 71, 93–100 (2018).

70. Prasad, R. et al. Specific interaction of DNA polymerase β and DNA ligase I in a multiprotein base excision repair complex from bovine testis. J. Biol. Chem., 271, 16000–16007 (1996).

71. Dimitriadis, E. K. et al. Thermodynamics of human DNA ligase I trimerization and association with DNA polymerase β. J. Biol. Chem., 273, 20540–20550 (1998).

72. Cotner-Gohara, E. et al. Human DNA ligase III recognizes DNA ends by dynamic switching between two DNA-bound states. Biochemistry 49, 6165–6176 (2010).

73. Taylor, R. M. et al. The DNA ligase III zinc finger stimulates binding to DNA secondary structure and promotes end joining. Nucleic Acids Res. 28, 3558–3563 (2000).

74. Cuneo, M. J. et al. The structural basis for partitioning of the XRCC1/DNA ligase III-a BRCT-mediated dimer complexes. Nucleic Acids Res. 39, 7816–7827 (2011).

75. Hammel, M. et al. An atypical BRCT-BRCT interaction with the XRCC1 scaffold protein compacts human DNA ligase IIIα within a flexible DNA repair complex. Nucleic Acids Res. 49, 306–321 (2021).

76. Conlin, M. P. et al. DNA ligase IV guides end-processing choice during nonhomologous end joining. Cell Rep. 20, 2810–2819 (2017).

77. Ochi, T., Gu, X. & Blundell, T. L. Structure of the catalytic region of DNA ligase IV in complex with an Artemis fragment sheds light on double-strand break repair. Structure 21, 672–679 (2013).

78. Ochi, T. et al. Structural insights into the role of domain flexibility in human DNA ligase IV. Structure 20, 1212–1222 (2012).

79. Kaminski, A. et al. Structures of DNA-bound human ligase IV catalytic core reveal insights into substrate binding and catalysis. Nat. Commun. 9, 2642 (2018).

80. Tomkinson, A. E., Tappe, N. J. & Friedberg, E. C. DNA Ligase I from *Saccharomyces cerevisiae*: Physical and biochemical characterization of the CDC9 gene product. Biochemistry 31, 11762–11771(1992).

81. Shi, K. et al. T4 DNA ligase structure reveals a prototypical ATP-dependent ligase with a unique mode of sliding clamp interaction. Nucleic Acids Res. 46, 10474–10488 (2018).

82. Bogenhagen, D. F. & Pinz, K. G. The action of DNA ligase at abasic sites in DNA. J. Biol. Chem. 273, 7888–7893 (1998).

83. Nishida, H., Kiyonari, S., Ishino, Y. & Morikawa, K. The closed structure of an archaeal DNA ligase from *Pyrococcus furiosus*. J. Mol. Biol. 360, 956–967 (2006).

84. Chen, Y. et al. Structure of the error-prone DNA ligase of African swine fever virus identifies critical active site residues. Nat. Commun. 10, 387 (2019).

85. Unciuleac, M., Goldgur, Y. & Shuman, S. Structures of ATP-bound DNA ligase D in a closed domain conformation reveal a network of amino acid and metal contacts to the ATP phosphates. J. Biol. Chem. 294, 5094–5104 (2019).

86. Shuman, S. & Lima, C. D. The polynucleotide ligase and RNA capping enzyme superfamily of covalent nucleotidyltransferases. Curr. Opin. Struct. Biol. 14, 757–764 (2004).

87. Pascal, J. M. DNA and RNA ligases: structural variations ands hared mechanisms. Curr. Opin. Struct. Biol. 18, 96–105 (2008).

88. Subramanya, H. S., Doherty, A. J., Ashford, S. R. & Wigley, D. B. Crystal structure of an ATP-dependent DNA ligase from bacteriophage T7. Cell 85, 607–615 (1996).

89. Odell, M., Sriskanda, V., Shuman, S. & Nikolov, D. Crystalstructure of eukaryotic DNA ligase–adenylate illuminates the mechanism of nick sensing and strand joining. Mol. Cell 6, 1183–1193 (2000).

90. Nair, P. A., Nandakumar, J., Smith, P., Odell, M., Lima, C. D. & Shuman, S. Structural basis for nick recognition by a minimal pluripotent DNA ligase. Nat. Struct. Mol. Biol. 14, 770–778 (2007).

91. Gong, C., Martins, A., Bongiorno, P., Glickman, M. & Shuman, S. Biochemical and genetic analysis of the four DNA ligases of mycobacteria. J. Biol. Chem. 279, 20594–20606 (2004).

92. Gong, C., Bongiorno, P., Martins, A., Stephanou, N. C., Zhu, H., Shuman, S. & Glickman, M. S. Mechanism of non-homologous end joining in mycobacteria: a low-fidelity repair system driven by Ku, ligase D and ligase C. Nat. Struct. Mol. Biol. 12, 304–312 (2005).

93. Akey, D., Martins, A., Aniukwu, J., Glickman, M. S., Shuman, S. & Berger, J. M. Crystal structure and nonhomologous end joining function of the ligase component of *Mycobacterium* DNA ligase D. J. Biol. Chem. 281, 13412–13423 (2006).

94. Williamson, A., Rothweiler, U. & Leiros, H. K. Enzyme-adenylate structure of a bacterial ATP-dependent DNA ligase with a minimized DNA-binding surface. Acta Crystallogr. D 70, 3043–3056 (2014).

95. Williamson, A., Grgic, M. & Leiros, H. S. DNA binding with a minimal scaffold: structure-function analysis of Lig E DNA ligases. Nucleic Acids Res. 46, 8616–8629 (2018).

96. Whitaker, A. M., Flynn, T. S. &. Freudenthal, B. D. Molecular snapshots of APE1 proofreading mismatches and removing DNA damage. Nat. Comm. 9, 399 (2019).

97. Andres, S. N. et al. Recognition and repair of chemically heterogeneous structures at DNA ends. Environ. Mol. Mutagen. 56, 1–21 (2015)

98. Guo, M. et al. Mechanism of genome instability mediated by human DNA polymerase mu misincorporation. Nat. Commun. 12, 3759 (2021).

99. Otwinowski, Z. & Minor, W. Processing of X-ray diffraction data collected in oscillation mode. Methods Enzymol. 276, 307–326 (1997).

100. McCoy, A. J. et al. Phaser crystallographic software. J. Appl. Crystallogr. 40, 658–674 (2007).

101. Emsley, P. et al. Features and development of Coot. Acta Crystallogr. D. Biol. Crystallogr. 66, 486–501 (2010).

102. Adams, P. D. et al. PHENIX: a comprehensive Python-based system for macromolecular structure solution. Acta Crystallogr. D. Biol. Crystallogr. 66, 213–221 (2010).

103. Murshudov, G. N. et al. REFMAC5 for the refinement of macromolecular crystal structures. Acta Crystallogr. D. Biol. Crystallogr. 67, 355–367 (2011).

