## Supplementary Data for BioRvix 2021-Caglayan for "Structures of LIG1 engaging with mutagenic mismatches inserted by polβ in base excision repair"

Supplementary Figures 1-11

Supplementary Schemes 1-3

Supplementary Tables 1-5

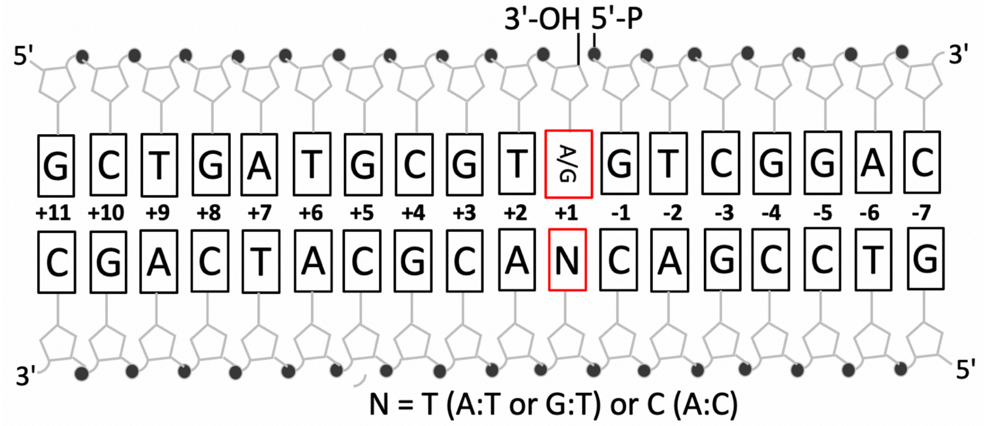


**Supplementary Figure 1.** Schematic view of upstream, downstream, and template oligonucleotide primers used to generate the nick DNA substrates with A:T, G:T, and A:C ends at 3'-strand for LIG1/nick DNA duplex crystallizations.

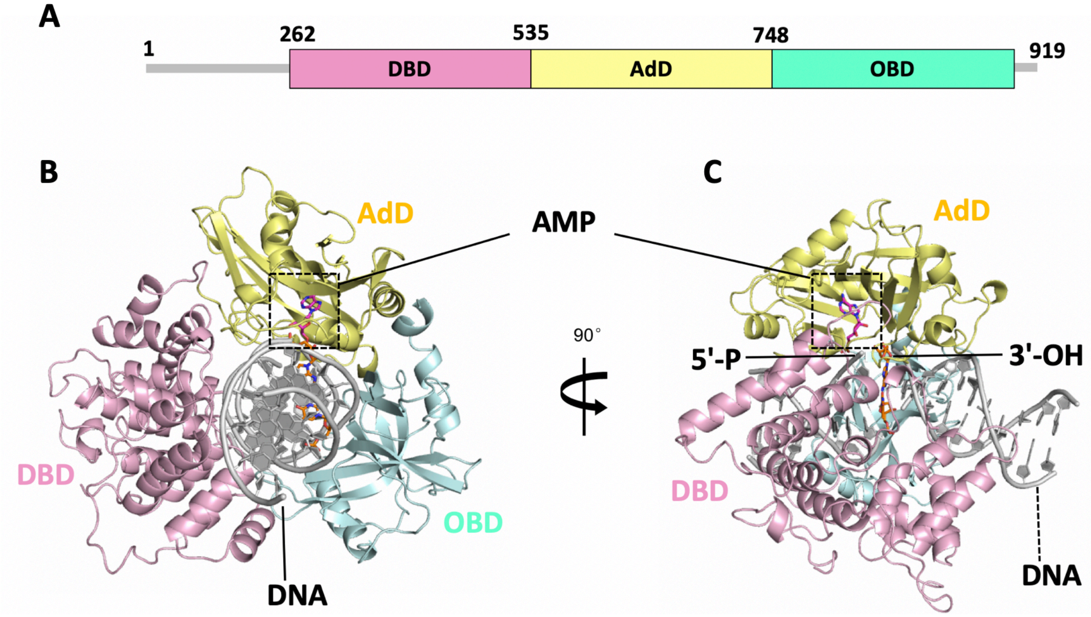


**Supplementary Figure 2.** LIG1 domain organization and structure. (**A**) Schematic view of LIG1 domain composition including the N-terminal domain (amino acids 1-261, gray), and the catalytic core (amino acids 262-919) consisting of the DNA-binding domain (DBD, pink), Adenylation domain (AdD, yellow), and OB-fold domain (OBD, green). (**B-C**) LIG1 (cartoon) domain encircling nick DNA is depicted in complex with AMP-DNA for A:T (B) and AMP-LIG1 for A:C (C) where AMP (orange) and DNA (grey) are shown as sticks at 3'-OH of nick (magenta).

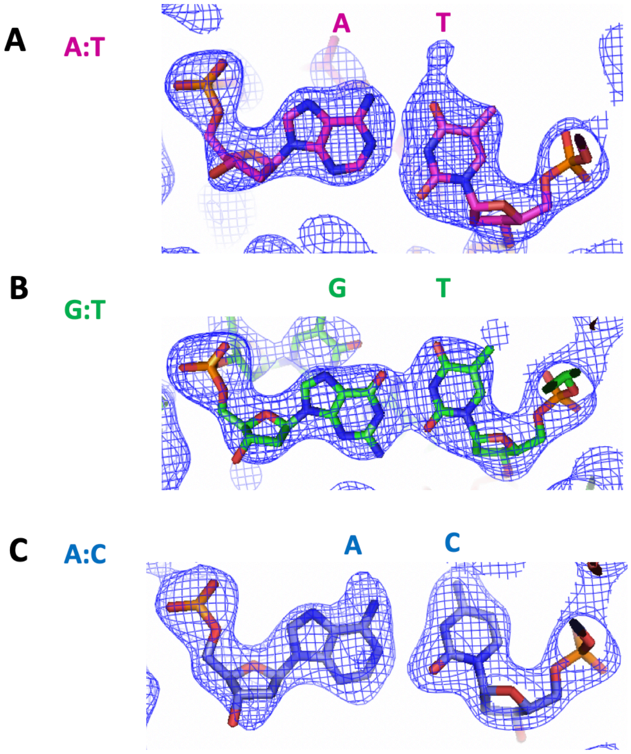


**Supplementary Figure 3.** 2Fo - Fc map density (1σ) 3'-OH terminus A:T (**A**), G:T (**B**) and A:C (**C**) are shown as blue. The structures of 3'-OH terminus at nick are shown as sticks for A:T (magenta), G:T (green) and A:C (blue).

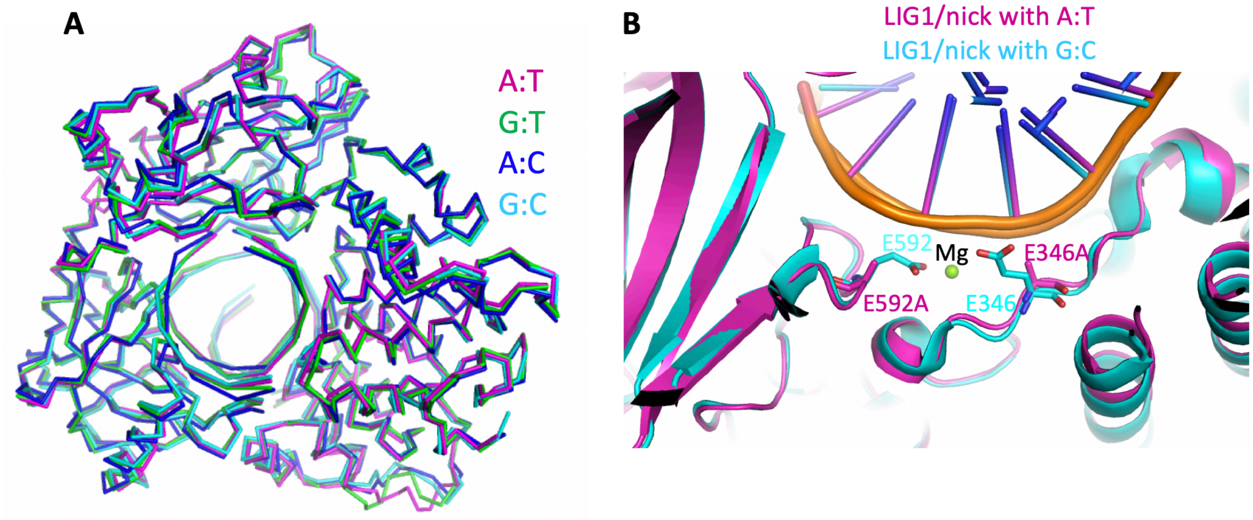


**Supplementary Figure 4. The comparison of LIG1/mismatched and matched nick DNA duplex structures.** (**A**) Overlay of wild-type LIG1/nick DNA duplexes with G:C (cyan) and EE/AA LIG1 nick DNA duplexes with A:T (magenta), G:T (green) and A:C (blue). All structures are shown as ribbons. (**B**) Superimposition of overall structures of wild-type LIG1 with G:C (PDB: 6P09, cyan) and EE/AA LIG1 with A:T (magenta) nick DNA duplexes is shown as cartoon and the amino acid residues E346 and E592 ($Mg^{HiFi}$ site) that are mutated in LIG1 EE/AA mutant are shown as sticks. $Mg^{2+}$ (green) is shown as sphere.

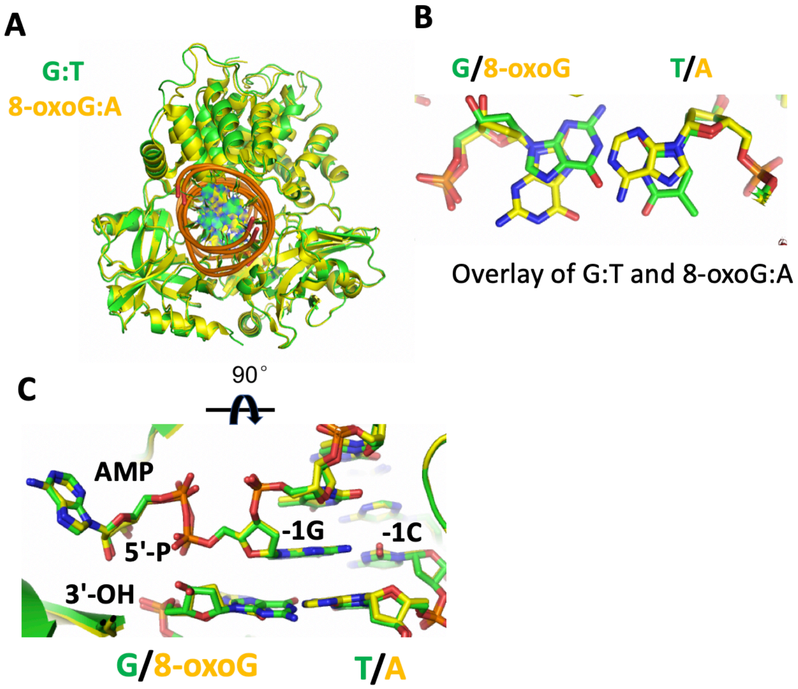


**Supplementary Figure 5. The comparison of LIG1/mismatched and damaged nick DNA duplex structures.** (**A**) Superimposition of overall structures of EE/AA LIG1 with G:T (green) and 8-oxoG:A (yellow, PDB:6POE) nick DNA duplexes are shown as cartoon. (**B,C**) overlay of G:T and 8-oxoG:A bound LIG1/nick DNA complexes (B) and the positions of 5'-P and 3'-OH ends at nick at the ligase active site (C).

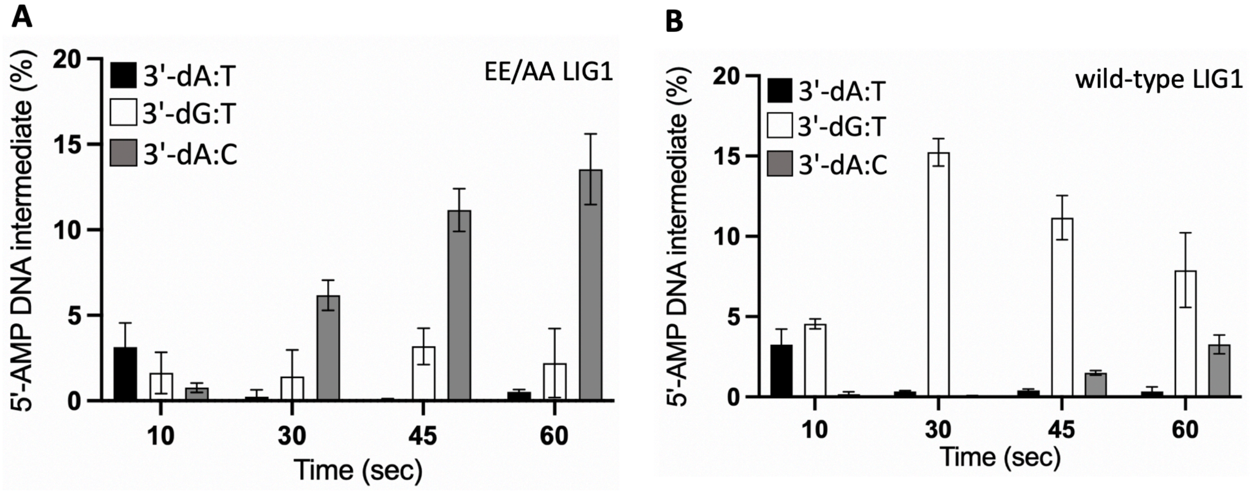


**Supplementary Figure 6. Ligation efficiency of nick repair intermediates with mismatches by LIG1.** (**A,B**) Graphs show time-dependent changes in the amount of DNA intermediates with 5'-AMP by EE/AA (A) and wild-type (B) of LIG1 for the nick DNA substrates with 3'-preinserted dA:T, dG:T, and dA:C. The data represent the average of three independent experiments ± SD. The gel images and ligation results are presented in Figure 6.

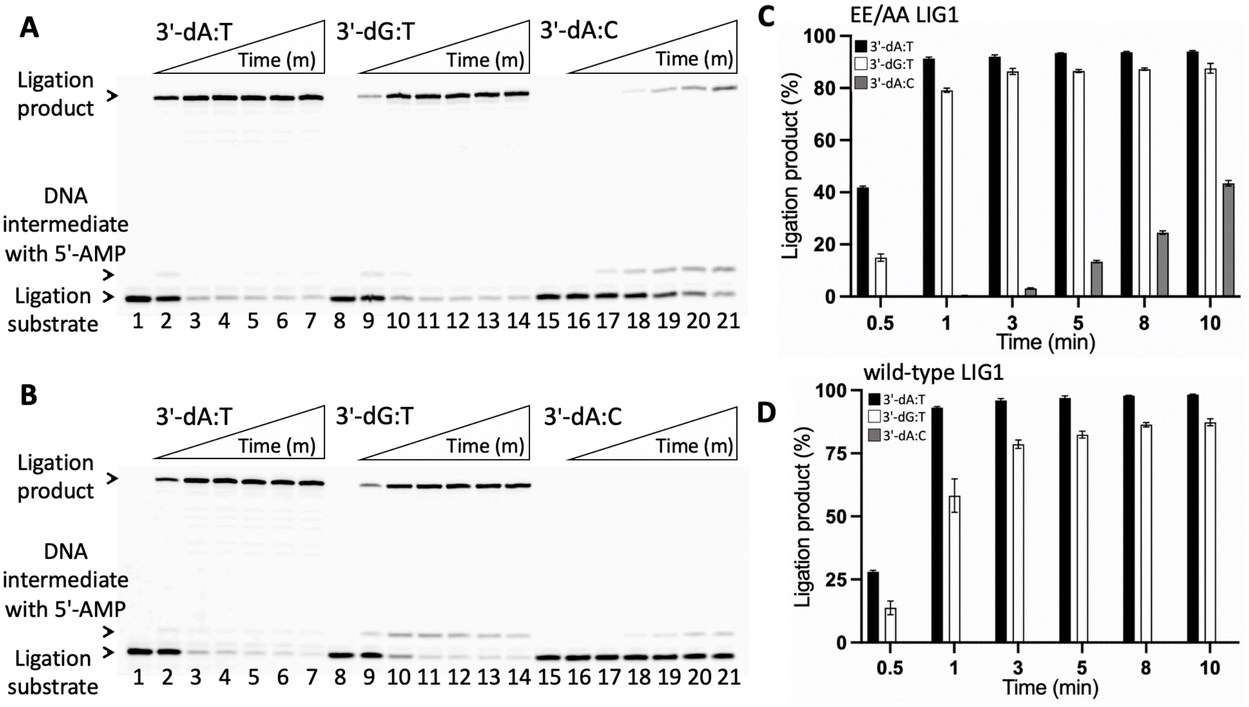


**Supplementary Figure 7. Ligation of nick repair intermediates with mismatches by LIG1.** (**A,B**) Lanes 1, 8, and 15 are the negative enzyme controls of the nick DNA substrates with 3'-preinserted dA:T, dG:T, and dA:C, respectively. Lanes 2-7, 9-14, and 16-21 are the reaction products for nick sealing of DNA substrates with 3'-preinserted dA:T, dG:T, and dA:C, respectively, by EE/AA (A) and wild-type (B) of LIG1, and correspond to time points of 0.5, 1, 3, 5, 8, and 10 min. (**C-D**) Graphs show time-dependent changes in the amount of ligation products. The data represent the average of three independent experiments ± SD.

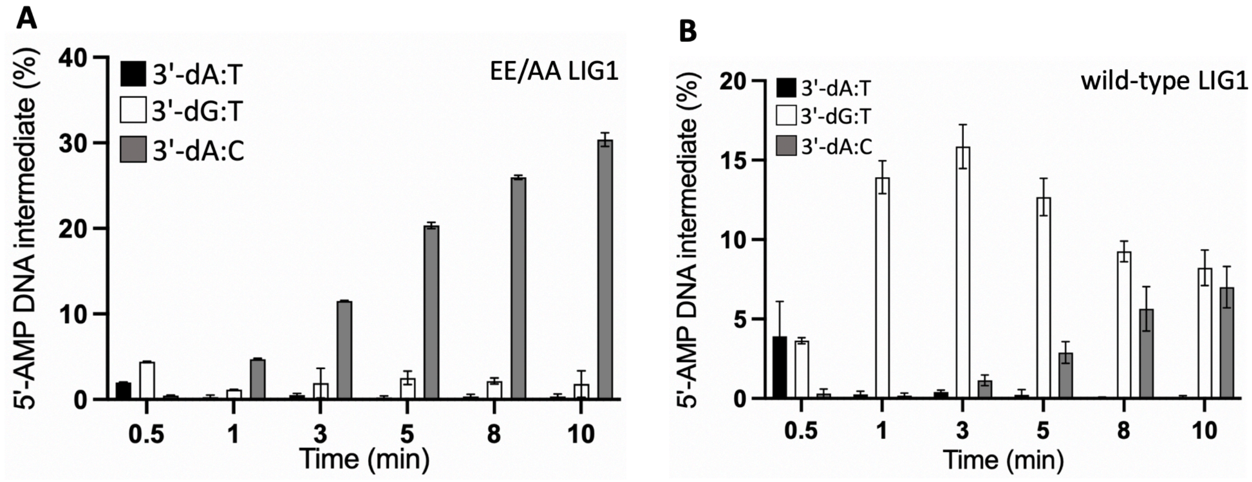


**Supplementary Figure 8. Ligation efficiency of nick repair intermediates with mismatches by LIG1.** (**A,B**) Graphs show time-dependent changes in the amount of DNA intermediates with 5'-AMP by EE/AA (A) and wild-type (B) of LIG1 for the nick DNA substrates with 3'-preinserted dA:T, dG:T, and dA:C. The data represent the average of three independent experiments ± SD. The gel images and ligation results are presented in Supplementary Figure 7.

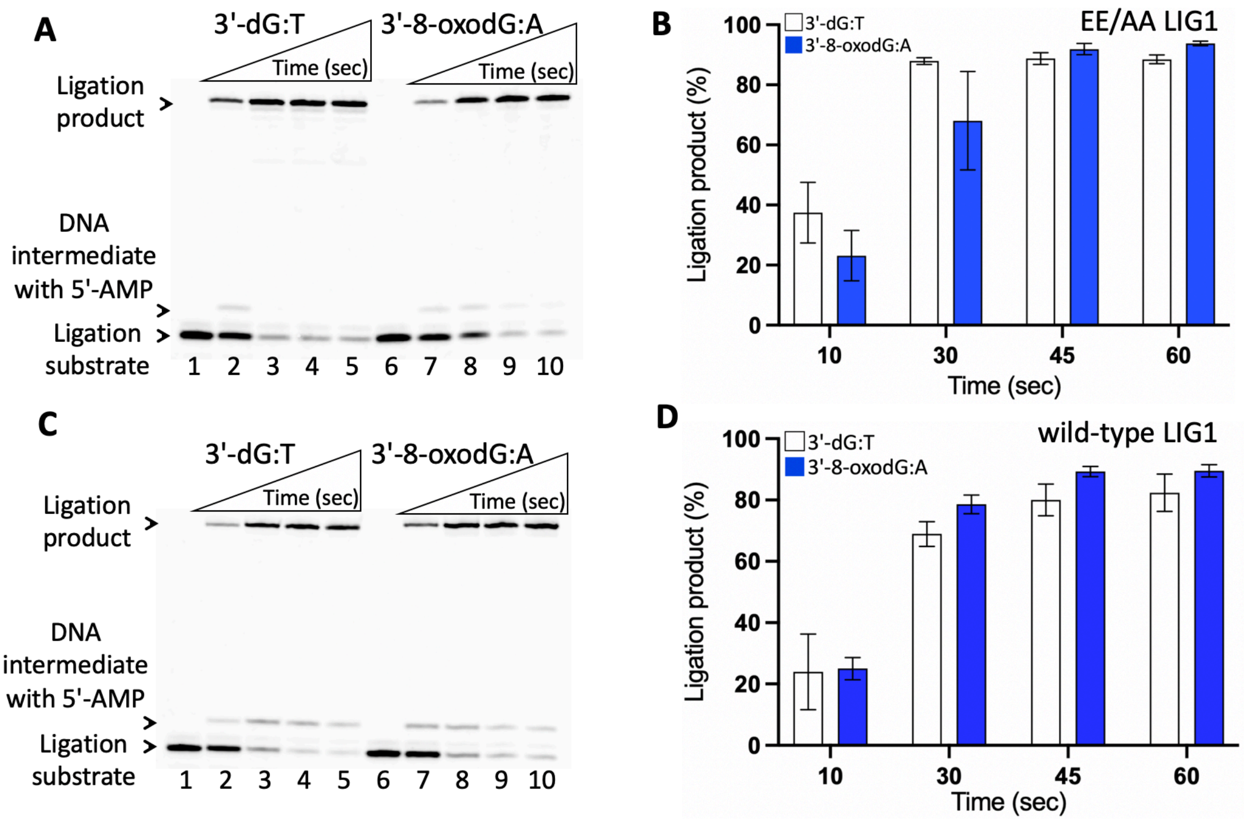


**Supplementary Figure 9. Ligation of nick repair intermediate with mismatch dG:T and damaged 8-oxodG:A by LIG1.** (**A,B**) Lanes 1 and 6 are the negative enzyme controls of the nick DNA substrates with 3'-preinserted dG:T and 8-oxodG:A, respectively. Lanes 2-5 and 7-10 are the reaction products for nick sealing of DNA substrates with 3'-preinserted dG:T and 8-oxodG:A by EE/AA (A) and wild-type (B) of LIG1, respectively, and correspond to time points of 10, 30, 45, and 60 sec. (**C-D**) Graphs show time-dependent changes in the amount of ligation products. The data represent the average of three independent experiments ± SD.

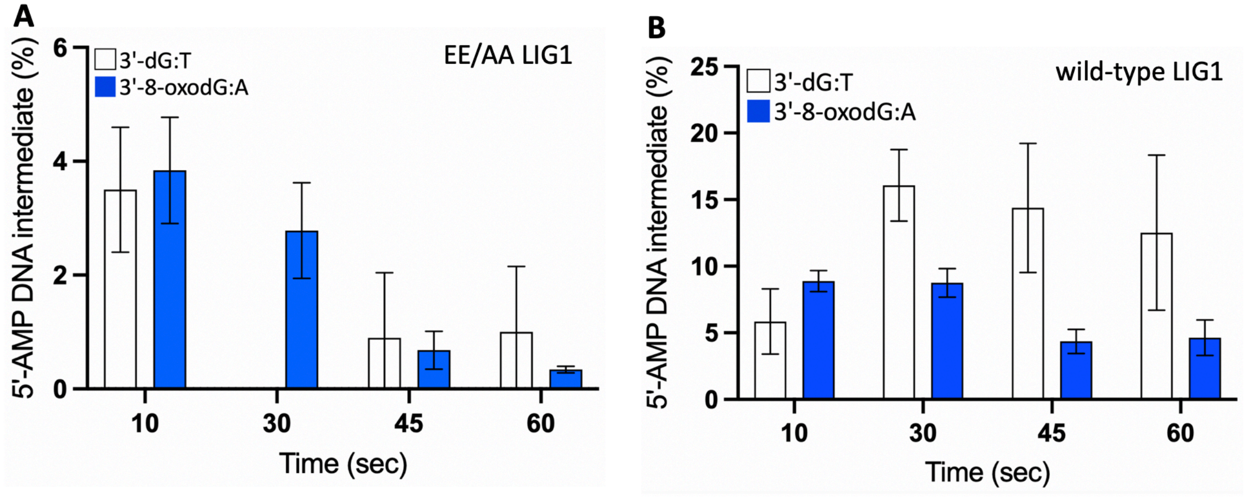


**Supplementary Figure 10. Ligation efficiency of nick repair intermediates with G:T mismatch and damaged 8-oxodG:A by LIG1.** (**A,B**) Graphs show time-dependent changes in the amount of DNA intermediates with 5'-AMP by EE/AA (A) and wild-type (B) of LIG1 for the nick DNA substrates with 3'-preinserted dG:T and 8-oxodG:A. The data represent the average of three independent experiments ± SD. The gel images and ligation results are presented in Supplementary Figure 9.

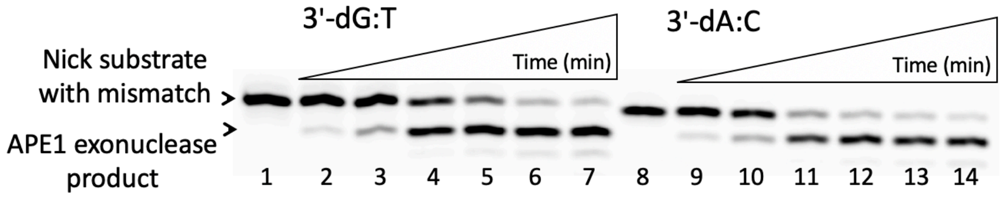


**Supplementary Figure 11. APE1 mismatch removal from nick repair intermediates with mismatches.** Lanes 1 and 8 are the negative enzyme controls of the nick DNA substrates with 3'-preinserted dG:T and dA:C, respectively. Lanes 2-7 and 9-14 are the reaction products of mismatch removal by APE1 from the nick DNA substrates with 3'-preinserted dG:T and dA:C, respectively, and correspond to time points of 0.5, 1, 2, 3, 4, and 5 min. Graph showing time-dependent changes in the amount of APE products is presented in Figure 8.

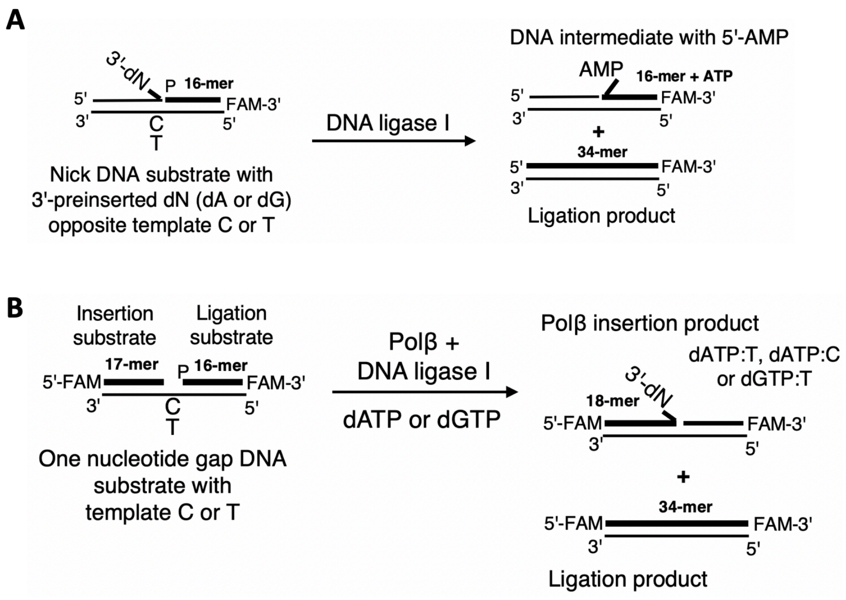


**Supplementary Scheme 1:** Illustrations of DNA repair assays used in this study. (**A**) Ligation assays were used to evaluate the substrate specificity of LIG1 for the nick DNA substrates including 3'-preinserted mismatches dG:T or dA:C and correctly base-paired dA:T. The reaction products include nick sealing or ligation product and DNA-AMP intermediates with 5'-adenylate (AMP). (**B**) Coupled assays were used to evaluate the ligation of polβ correct (dGTP:C) or mismatch (dGTP:T or dATP:C) insertions in the reaction mixture including polβ and LIG1 using one nucleotide gap DNA substrates with template base C or T. The reaction products include polβ nucleotide insertion and ligation products.

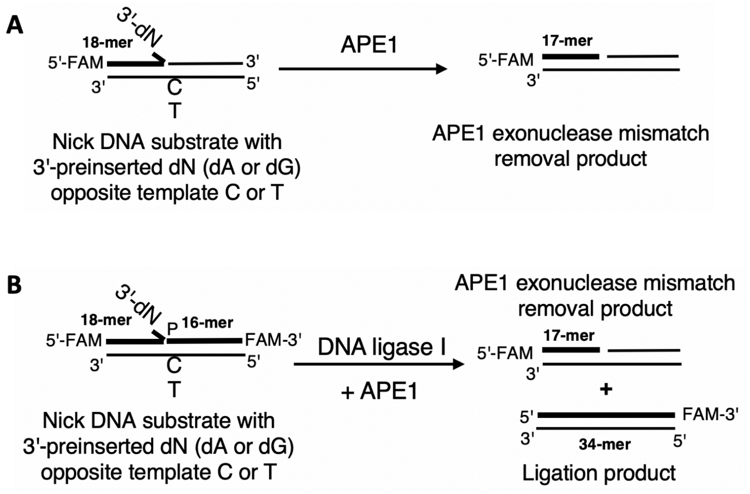


**Supplementary Scheme 2:** Illustrations of APE1 repair assays used in this study. (**A**) APE1 exonuclease activity assays were used to evaluate its proofreading role for the removal of a mismatched base from the nick DNA substrates including 3'-preinserted mismatches dG:T or dA:C. (**B**) Repair assays were used to evaluate the ligation and mismatch removal in the reaction mixture including APE1 and LIG1 using the nick DNA substrates including 3'-preinserted mismatches dG:T or dA:C. The reaction products include APE1 exonuclease removal and ligation products.

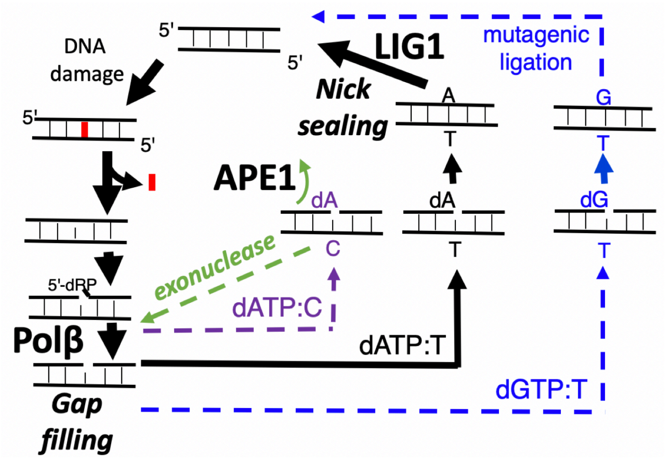


**Supplementary Scheme 3:** Illustration of the downstream steps involving polβ gap filling and subsequent nick sealing in the BER pathway. The model shows the efficient nick sealing of nick repair intermediate after polβ correct dATP:T insertion. Polβ dGTP insertion opposite T results in the mutagenic ligation of nick repair intermediate with G:T mismatch by LIG1. In contrast, polβ dATP mismatch insertion opposite C cannot be ligated during the subsequent ligation step, which could serve as a structural fidelity checkpoint and a signal for proofreading enzyme such as APE1 for a mismatched base removal.

| DNA substrates | Sequence |
|---|---|
| Nick DNA with preinserted 3'-dG:T | 5'-CATGGGCGGCATGAACCGGAGGCCCATCCTCACC-3'-FAM<br>3'-GTACCCGCCGTACTTGGTCTCCGGGTAGGAGTGG-5' |
| Nick DNA with preinserted 3'-dA:C | 5'-CATGGGCGGCATGAACCAGAGGCCCATCCTCACC-3'-FAM<br>3'-GTACCCGCCGTACTTGGCCTCCGGGTAGGAGTGG-5' |
| Nick DNA with preinserted 3'-8-oxodG:A | 5'-CATGGGCGGCATGAACCXGAGGCCCATCCTCACC-3'-FAM<br>3'-GTACCCGCCGTACTTGGACTCCGGGTAGGAGTGG-5' |

**Supplementary Table 1**. The nick DNA substrates used in ligation assays. FAM denotes a flourescence tag and is located at 3'-end of DNA substrates. The base at template base position is underlined.

| DNA substrates | Sequence |
|---|---|
| One nucleotide gap DNA with template base C | FAM-5’-CATGGGCGGCATGAACC GAGGCCCATCCTCACC-3’-FAM<br>3’-GTACCCGCCGTACTTGGCCTCCGGGTAGGAGTGG-5’ |
| One nucleotide gap DNA with template base T | FAM-5’-CATGGGCGGCATGAACC GAGGCCCATCCTCACC-3’-FAM<br>3’-GTACCCGCCGTACTTGGTCTCCGGGTAGGAGTGG-5’ |

**Supplementary Table 2**. The one nucleotide gap DNA substrates used in repair assays to test ligation of polβ nucleotide insertion products by LIG1 on coupled repair assays. FAM denotes a flourescence tag and is located at 5'- and 3'-ends of DNA substrates. The base at template base position is underlined.

| DNA substrates | Sequence |
|---|---|
| Nick DNA with preinserted 3'-dG:T | FAM-5’-CATGGGCGGCATGAACCGGAGGCCCATCCTCACC-3’<br>3’-GTACCCGCCGTACTTGGTCTCCGGGTAGGAGTGG-5’ |
| Nick DNA with preinserted 3'-dA:C | FAM-5’-CATGGGCGGCATGAACCAGAGGCCCATCCTCACC-3’<br>3’-GTACCCGCCGTACTTGGCCTCCGGGTAGGAGTGG-5’ |

**Supplementary Table 3**. The nick DNA substrates used in APE1 exonuclease assays. FAM denotes a flourescence tag and is located at 5'-end of DNA substrates. The base at template base position is underlined and P is phosphate group.

| DNA substrates | Sequence |
|---|---|
| Nick DNA with preinserted 3'-dG:T | FAM-5’-CATGGGCGGCATGAACCGGAGGCCCATCCTCACC-3’-FAM<br>3’-GTACCCGCCGTACTTGGTCTCCGGGTAGGAGTGG-5’ |
| Nick DNA with preinserted 3'-dA:C | FAM-5’-CATGGGCGGCATGAACCAGAGGCCCATCCTCACC-3’-FAM<br>3’-GTACCCGCCGTACTTGGCCTCCGGGTAGGAGTGG-5’ |

**Supplementary Table 4**. The nick DNA substrates used in the coupled repair assays for APE1 exonuclease mismatch removal and ligation by LIG1. FAM denotes a flourescence tag and is located at 3'- and 5'-end of DNA substrates. The base at template base position is underlined.

| Oligonucleotide | Primer | Sequence (5'-3') |
|---|---|---|
| 1 | Template T | GTCCGACTACGCATCAGC |
| 2 | Template C | GTCCGACCACGCATCAGC |
| 3 | Upstream A | GCTGATGCGTA |
| 4 | Upstream G | GCTGATGCGTG |
| 5 | Downstream | P-GTCGGAC |

**Supplementary Table 5.** The oligonucleotide primers used for LIG1 crystallization. Oligonucleotides 1, 3, and 5 were used to prepare the nick DNA substrate with 3'-dA:T. Oligonucleotides 1, 4, and 5 were used to prepare the nick DNA substrate with 3'-dG:T. Oligonucleotides 2, 3, and 5 were used to prepare the nick DNA substrate with 3'-dA:C.
